# *Caenorhabditis elegans* ETR-1/CELF has broad effects on the muscle cell transcriptome, including genes that regulate translation and neuroblast migration

**DOI:** 10.1101/2021.06.08.447597

**Authors:** Matthew E. Ochs, Rebecca M. McWhirter, Robert L. Unckless, David M. Miller, Erik A. Lundquist

## Abstract

Migration of neuroblasts and neurons from their birthplace is central to the formation of neural circuits and networks. ETR-1 is the *Caenorhabditis elegans* homolog of the CELF1 (<u>C</u>UGBP, <u>EL</u>AV-like family 1) RNA-processing factor involved in neuromuscular disorders. *etr-1* regulates body wall muscle differentiation. Our previous work showed that *etr-1* in muscle has a non-autonomous role in neuronal migration, suggesting that ETR-1 is involved in the production of a signal emanating from body wall muscle that controls neuroblast migration and that interacts with Wnt signaling. *etr-1* is extensively alternatively-spliced, and we identified the viable *etr-1(lq61)* mutant, caused by a stop codon in alternatively-spliced exon 8 and only affecting *etr-1* isoforms containing exon 8. We took advantage of viable *etr-1(lq61)* to identify potential RNA targets of ETR-1 in body wall muscle using a combination of fluorescence activated cell sorting (FACS) of body wall muscles from wild-type and *etr-1(lq61)* and subsequent RNA-seq. This analysis revealed genes whose splicing and transcript levels were controlled by ETR-1 exon 8 isoforms, and represented a broad spectrum of genes involved in muscle differentiation, myofilament lattice structure, and physiology. Genes with transcripts underrepresented in *etr-1(lq61)* included those involved in ribosome function and translation, similar to potential *CELF1* targets identified in chick cardiomyocytes. This suggests that at least some targets of ETR-1 might be conserved in vertebrates, and that ETR-1 might generally stimulate translation in muscles. As proof-of-principle, a functional analysis of a subset of ETR-1 targets revealed genes involved in AQR and PQR neuronal migration. One such gene, *lev-11/tropomyosin*, requires ETR-1 for alternative splicing, and another, *unc-52/perlecan*, requires ETR-1 for the production of long isoforms containing 3’ exons. In sum, these studies identified gene targets of ETR-1/CELF1 in muscles, which included genes involved in muscle development and physiology, and genes with novel roles in neuronal migration.

## Introduction

Migration of neuroblasts and neurons is a key developmental process in the formation of neural circuits and networks. The CELF (<u>C</u>UGBP, <u>EL</u>AV-like family) class of RNA-binding proteins is implicated in a wide variety of neuromuscular and neurodegenerative disorders, including Myotonic Dystrophy type I (DMI) (Savkur *et al*. 2001; Timchenko *et al*. 2001; Timchenko *et al*. 2004; Ho *et al*. 2005; Kuyumcu-Martinez *et al*. 2007; Schoser and Timchenko 2010; Berger and Ladd 2012), the cardiac syndrome arrythmogenic right ventricular dysplasia (Li *et al*. 2001), Alzheimer’s disease (Wijsman *et al*. 2011), spinocerebellar ataxia type 8, and possibly fragile X syndrome (Sofola *et al*. 2007; Daughters *et al*. 2009). CELF proteins control mRNA processing, including alternative splicing (Ladd *et al*. 2005; Kalsotra *et al*. 2008; Terenzi *et al*. 2009), and regulation of translation (Kim *et al*. 2015), and mRNA transcript stability (Dasgupta and Ladd 2012; Blech-Hermoni *et al*. 2016). CELF protein structure is characterized by three RNA-recognition motifs (RRM) with a non-conserved region between RRM2 and RRM3, an RRM organization conserved across CELF protein family members (Dasgupta and Ladd 2012).

Vertebrate genomes encode up to six CELF molecules (Good *et al*. 2000), CELF1-6. In *C. elegans,* there are two *CELF* genes, *etr-1* (most similar to *CELF1-2*) (Milne and Hodgkin 1999; Ochs *et al*. 2020), and *unc-75* (most similar to *CELF3-6)* (Loria *et al*. 2003). *unc-75* controls alternative splicing events predominantly in the nervous system (Loria *et al*. 2003; Kuroyanagi *et al*. 2013a; Kuroyanagi *et al*. 2013b; Norris *et al*. 2014; Chen *et al*. 2016). ETR-1 is involved in muscle development, as knockdown of ETR-1 results in severe muscle disorganization and embryonic lethality (Milne and Hodgkin 1999). ETR-1 also controls cell corpse engulfment in the germline (Boateng and Allen 2018), and influences neuronal migration non-autonomously from body wall muscle (Ochs *et al*. 2020).

ETR-1 acts in muscle to guide the long-range migration of the Q neuroblast descendants in *C. elegans* (Ochs *et al*. 2020). The Q neuroblasts, QR and QL, are bilaterally symmetrical cells that undergo similar divisions and stereotypical migrations (reviewed in (Middelkoop and Korswagen 2014)). QR is born on the right side of the animal and QR descendants migrate anteriorly. QL is born on the left side of the animal and QL descendants migrate posteriorly. Both QR and QL produce three functional neurons, and two apoptotic bodies. QR produces AQR, AVM and SDQR, with AQR migrating the furthest, residing just posterior to the posterior pharyngeal bulb in the anterior deirid ganglion. QL produces PQR, PVM and SDQL, with PQR migrating the furthest, residing posterior to the anus in the phasmid ganglion (Sulston and Horvitz 1977; Chalfie and Sulston 1981; Chapman *et al*. 2008). Due to the stereotypical migrations, the Q neuroblasts are a powerful system to identify migration defects and study the genetic mechanisms controlling migration. Initial QL and QR migrations are controlled by interactions between three receptor molecules UNC-40/DCC, PTP-3/LAR and MIG-21, and the Fat-like Cadherins CDH-3 and CDH-4 (Honigberg and Kenyon 2000; Middelkoop *et al*. 2012; Sundararajan and Lundquist 2012; Sundararajan *et al*. 2014; Ebbing *et al*. 2019). Migration of the Q descendants is controlled by Wnt signaling along the anterior-posterior body axis (Zinovyeva *et al*. 2008; Harterink *et al*. 2011; Josephson *et al*. 2016a). Previous studies have implicated body wall muscle cells as sources of migration factors for the Q neuroblast descendants (Josephson *et al*. 2016b; Josephson *et al*. 2017).

The *etr-1(lq61)* mutation was isolated in a forward genetic screen for AQR and PQR migration defects (Ochs *et al*. 2020). *lq61* introduces a premature stop codon in alternatively-spliced exon 8, which is present in a subset of *etr-1* isoforms. Total knockdown of *etr-1* resulted in embryonic lethality with muscle defects (Milne and Hodgkin 1999), yet *etr-1(lq61)* animals are viable and fertile, consistent with *lq61* being a hypomorphic mutation. *etr-1* is expressed in all cells of the embryo (Boateng and Allen 2018), but acts in the muscle cells in a non-autonomous manner to control AQR and PQR migrations (Ochs *et al*. 2020). Furthermore, *etr-1(lq61)* interacts genetically with *Wnt* mutations in AQR and PQR migration.

As *etr-1(lq61)* is a viable and fertile mutation, it presented a unique opportunity to identify gene targets of a CELF1/2 family member in muscles, and to define target genes that contribute to the non-autonomous control of AQR and PQR migration by ETR-1. We used fluorescence activated cell sorting of *C. elegans* body wall muscle cells from wild-type and *etr-1(lq61)* mutants, combined with RNA-seq, to define muscle-expressed genes with alternative exon usage and transcript accumulation and in *etr-1(lq61)* mutants. This analysis revealed genes involved in myofilament lattice assembly and adhesion, and muscle physiology. Genes with underrepresented transcripts in *etr-1(lq61)* were involved in translation and ribosome function. As proof of principle, a pilot functional screen identified new genes for AQR and PQR migration, including *unc-52/perlecan* and *lev-11/tropomyosin*. ETR-1 targets, including *lev-11/tropomyosin* and genes involved in translational and ribosome function, were also identified in vertebrates (DASGUPTA AND LADD 2012; BLECH-HERMONI *et al*. 2016), suggesting a deep evolutionary conservation of CELF targets and potentially conserved molecular mechanisms of CELF1/2 function from *C. elegans* to vertebrates.

## Results

### Fluorescent-activated cell sorting of muscle cells and RNA seq

*myo-3::gfp*-expressing body wall muscle cells from synchronized early L1 larvae were isolated by FACS as described in Material and Methods and in (Spencer *et al*. 2014; Taylor *et al*. 2020) (Figure 1). Muscles were isolated from the wild-type (N2) strain and *etr-1(lq61)* mutants. Three biological replicates for each genotype were isolated. RNA was also isolated from triplicate samples of non-dissociated L1 larvae for the all-cell control group.

**Figure 1.**
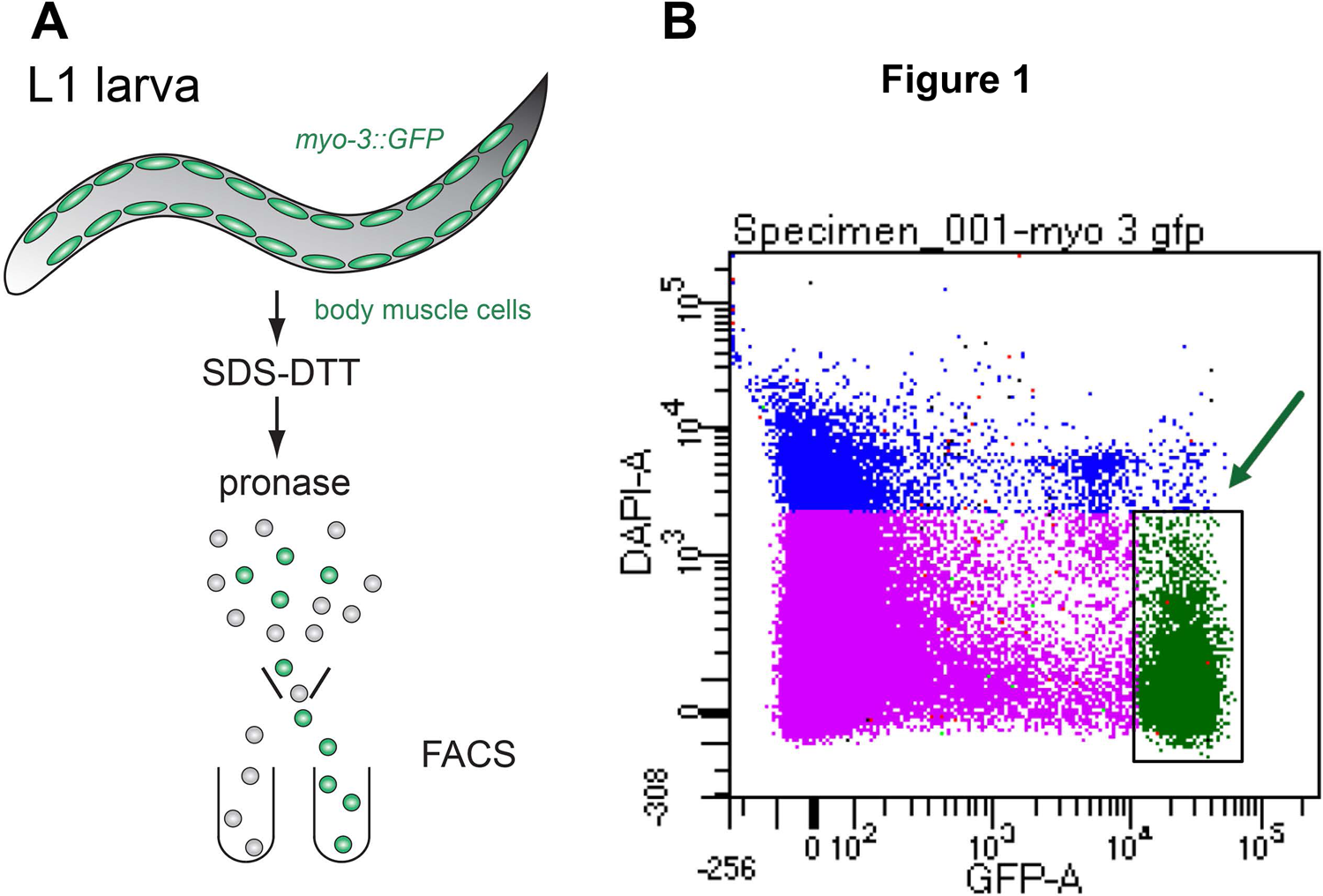
FACS isolation of body muscle cells from wild type and *etr-1(lq61)* mutant L1 larvae. A) Synchronized L1-stage larvae were dissociated by successive treatments with SDS-DTT and pronase to release *myo-3::gfp* labeled body muscle cells for isolation by FACS. B) FACS scatter plot. Viable and brightly labeled myo-3::GFP marked cells (arrow) were captured for RNA extraction. Damaged cells were excluded by DAPI staining.

A total of twelve RNA-seq libraries were constructed, representing three biological replicates of both N2 and *etr-1(lq61)* with both sorted muscle cells and whole L1 larval stage cells (see Materials and Methods). Paired-end 150-bp reads were generated from each of the twelve samples using the Illumina Nextseq550 platform FASTQ files can be accessed in the Sequence Read Archive, Project number PRJNA733501

### A muscle cell transcriptome defined by RNA-seq

We used DEseq2 to identify genes with significant differential expression in muscle cells compared to all L1 stage cells (Supplemental File 1). There were 3,718 protein-coding genes with significantly higher expression in muscle cells compared to all cells (log2-fold change ≥ 0.5849 (1.5x); *q* ≤ 0.05), including many canonical muscle structure and function genes previously shown to be expressed in muscle (e.g. *unc-15/paramyosin, unc-54/myosin, unc-95/paxillin,* and the myofilament structure *pat* genes (Fox *et al*. 2007). DEseq2 also identified *etr-1* as being more highly expressed in muscle, with multiple exons showing significantly increased expression (Figure 2A). The *etr-1* locus is extensively alternatively spliced (BOATENG *et al*. 2017; Ochs *et al*. 2020), and isoforms with exon 8 are required in muscles for Q neuroblast descendant migration (Ochs *et al*. 2020). Exon 8 expression was increased in muscles compared to all cells (Figure 2A). Expression of 8,763 protein coding genes was significantly reduced in muscle cells compared to all cells (Supplemental File 1). Differential expression of non-coding RNAs and pseudogenes is presented in Supplemental File 1.

**Figure 2.**
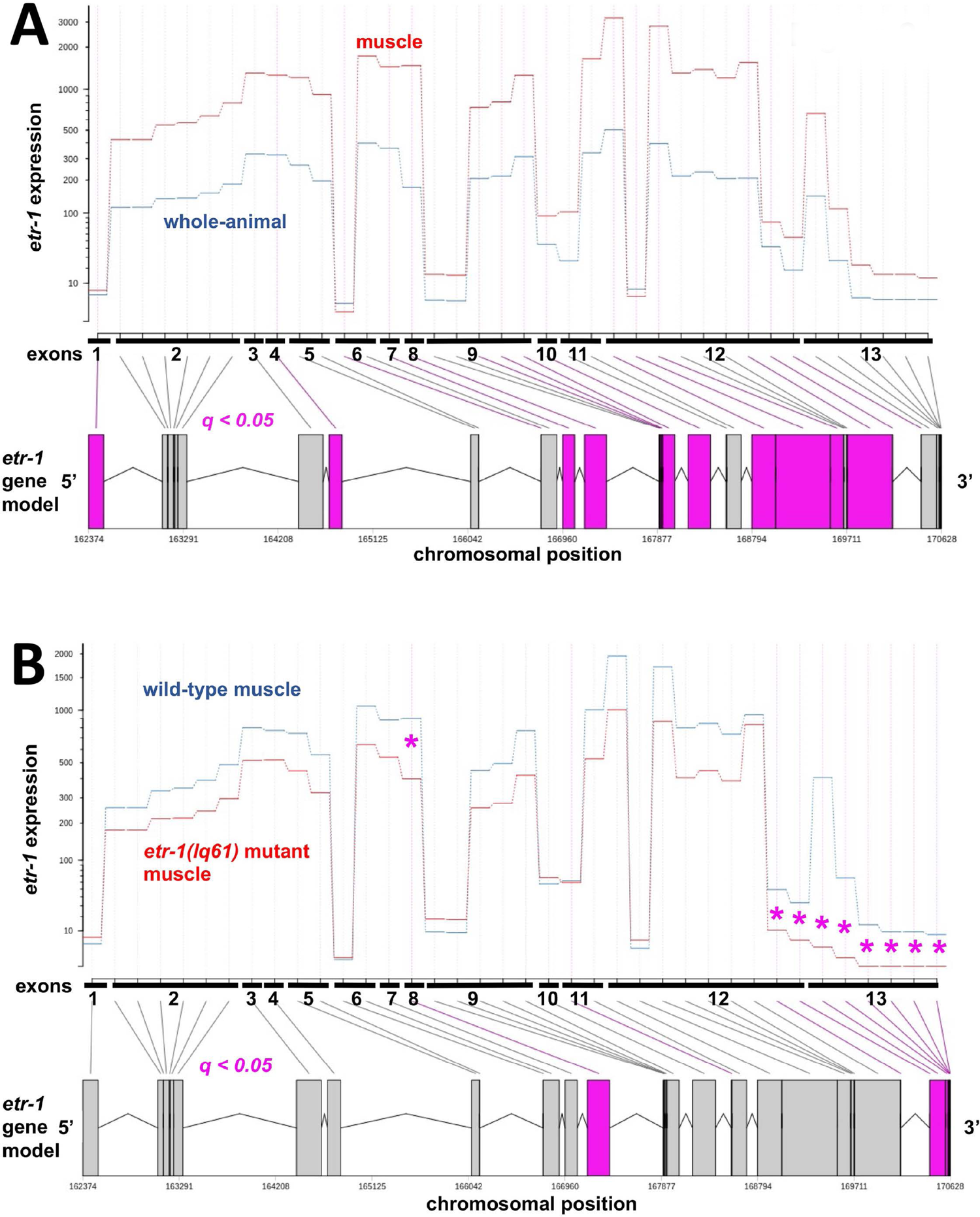
Output of DEXSeq showing differential exon usage. A) Exon usage of *etr-1* comparing wild-type muscle to wild-type whole animal. The red line represents muscle expression, the blue line represents whole animal expression. Exons that are significantly different (*q* < 0.05) are in purple. *etr-1* exons were overrepresented in muscle cells, including exon 8 harboring the *lq61* mutation. B) Exon usage of *etr-1* comparing *etr-1(lq61)* muscle cells to wild-type muscle cells. The red line represents *etr-1(lq61)* expression and the blue line represents wild-type expression. *etr-1* exons generally were underrepresented in *etr-1(lq61)*, with exon 8 and the 3’ exon significantly so (purple).

### Genes with exon usage affected by *etr-1(lq61)* encode molecules involved in myofilament lattice structure and attachment, and muscle physiology

CELF family proteins are known to regulate splicing (Ladd *et al*. 2001; Ladd *et al*. 2005), and we endeavored to determine the effects of ETR-1/CELF on the muscle transcriptome, including splicing. We compared exon representation across the genome in wild-type and *etr-1(lq61)* mutant muscle cells using the Bioconductor package DEXseq (Anders *et al*. 2013) (see Materials and Methods). Across the genome, there were 242 protein-coding genes and seven non-coding RNA genes with at least one exon significantly differentially represented in *etr-1(lq61)* muscle compared to wild-type (*q* ≤ 0.05) (Supplemental File 2).

The *etr-1(lq61)* mutation is a premature stop codon in alternatively-spliced exon 8 (Ochs *et al*. 2020). Exon 8 was significantly underrepresented in *etr-1(lq61)* muscle compared to wild-type (Figure 2B), suggesting that transcripts containing exon 8 are reduced in *etr-1(lq61)* muscle cells, as predicted. Exon 13 was also significantly underrepresented in *etr-1(lq61).* Possibly, transcripts with exon 8 might preferentially contain exon 13. Alternatively, ETR-1 containing exon 8 might be involved in the regulation of processing of *etr-1* exon 13 We used the Database for Annotation, Visualization and Integrated Discovery (DAVID) (Huang da *et al*. 2009b; Huang da *et al*. 2009a) to perform a gene ontology term (GO term) analysis on this gene set that showed differential exon representation in *etr-1(lq61)* mutant muscle compared to wild-type muscle. (see Materials and methods). We analyzed GO terms for the three categories: biological process (BP), cellular component (CC) and molecular function (MF) (Supplemental File 3). The six most significantly enriched GO terms in each category are shown in Figure 3. These include GO terms associated with myofilament lattice formation and function (e.g. striated muscle myosin thick filament assembly, locomotion, M band, striated muscle thin filament assembly, striated muscle dense body, I band, actin filament, and actin filament binding). Also included are muscle physiology GO terms (e.g. voltage gated ion channel activity, calcium ion binding, voltage gated potassium channel activity, and kinase activity). These are consistent with the previously-reported effects of *etr-1* RNAi knockdown on muscle development and attachment (Milne and Hodgkin 1999). GO terms of apoptotic process and reproduction are also enriched, consistent with a known role of *etr-1* in germline development and engulfment of germ cell apoptotic corpses (Boateng *et al*. 2017).

**Figure 3.**
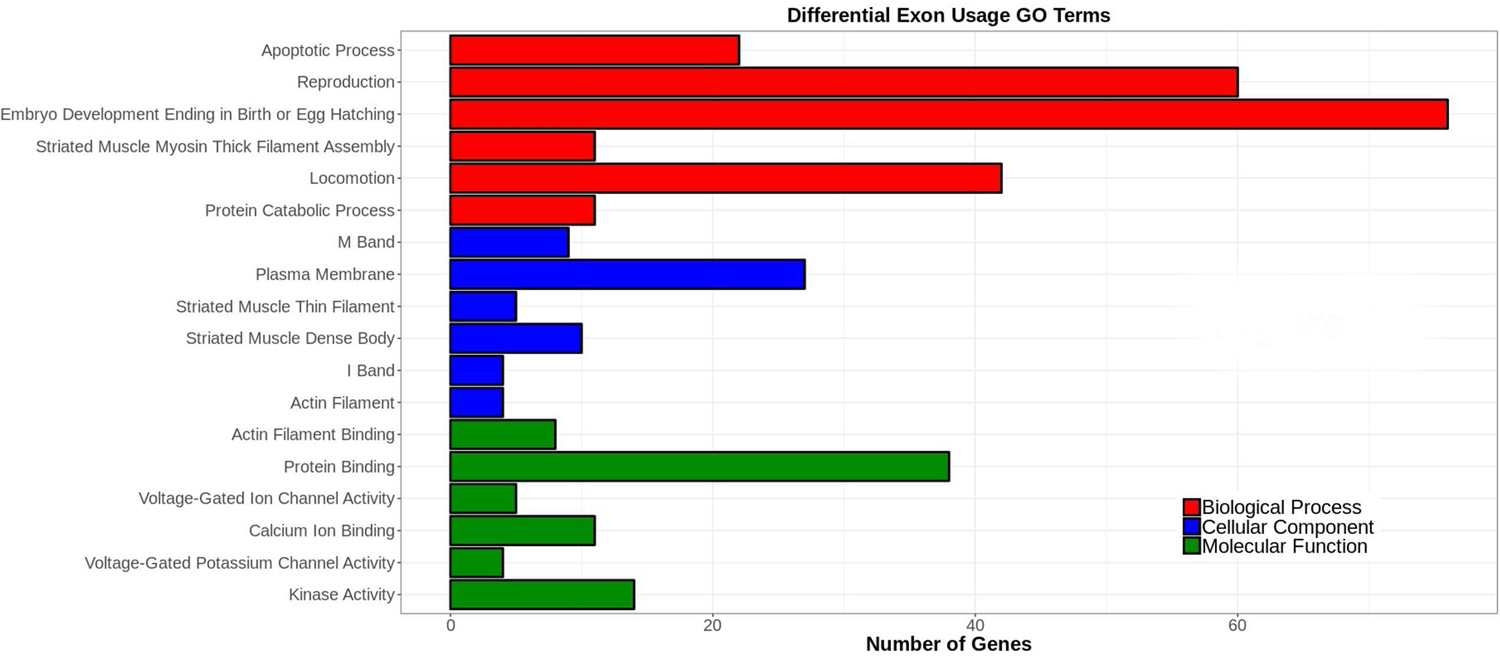
Gene ontology (GO) term analysis of genes with differential exon representation in *etr-1(lq61)*. Analysis was done using the Database for Annotation, Visualization and Integrated Discovery (DAVID) (see Materials and Methods). GO terms are along the Y axis, and the number of genes associated with each GO term is along the X axis. The six most significant GO terms (*q* ≤ 0.05) for three categories (biological process, cellular component, and molecular function) are shown. The entire GO term analysis is in Supplemental File 3.

### Genes with transcript accumulation affected by *etr-1(lq61)* encode molecules involved in translation and ribosome function

The CELF-family proteins control transcript stability (Ladd *et al*. 2001; Ladd *et al*. 2005; Kalsotra *et al*. 2008). We used stringTie and DEseq2 to identify transcripts with differential accumulation in wild-type versus *etr-1(lq61)* muscle cells (see Materials and Methods). We identified transcripts that were differentially represented with a log_2_ fold change ≥ 1 (2x) and a false discovery rate less than 0.05 (*q* ≤ 0.05), to increase stringency given the large number of genes returned in this analysis (Supplemental File 4).

We identified 1180 transcripts representing 971 loci with altered accumulation in *etr-1(lq61)* (Figure 4A), including coding and non-coding RNAs (Supplemental File 4). 506 loci had transcripts that were overrepresented, and 414 loci had transcripts that were underrepresented in *etr-1(lq61)* muscle cells (Figure 4B). There were 51 loci in which some transcripts were overrepresented and some underrepresented (Figure 4B and Table 1).

**Figure 4.**
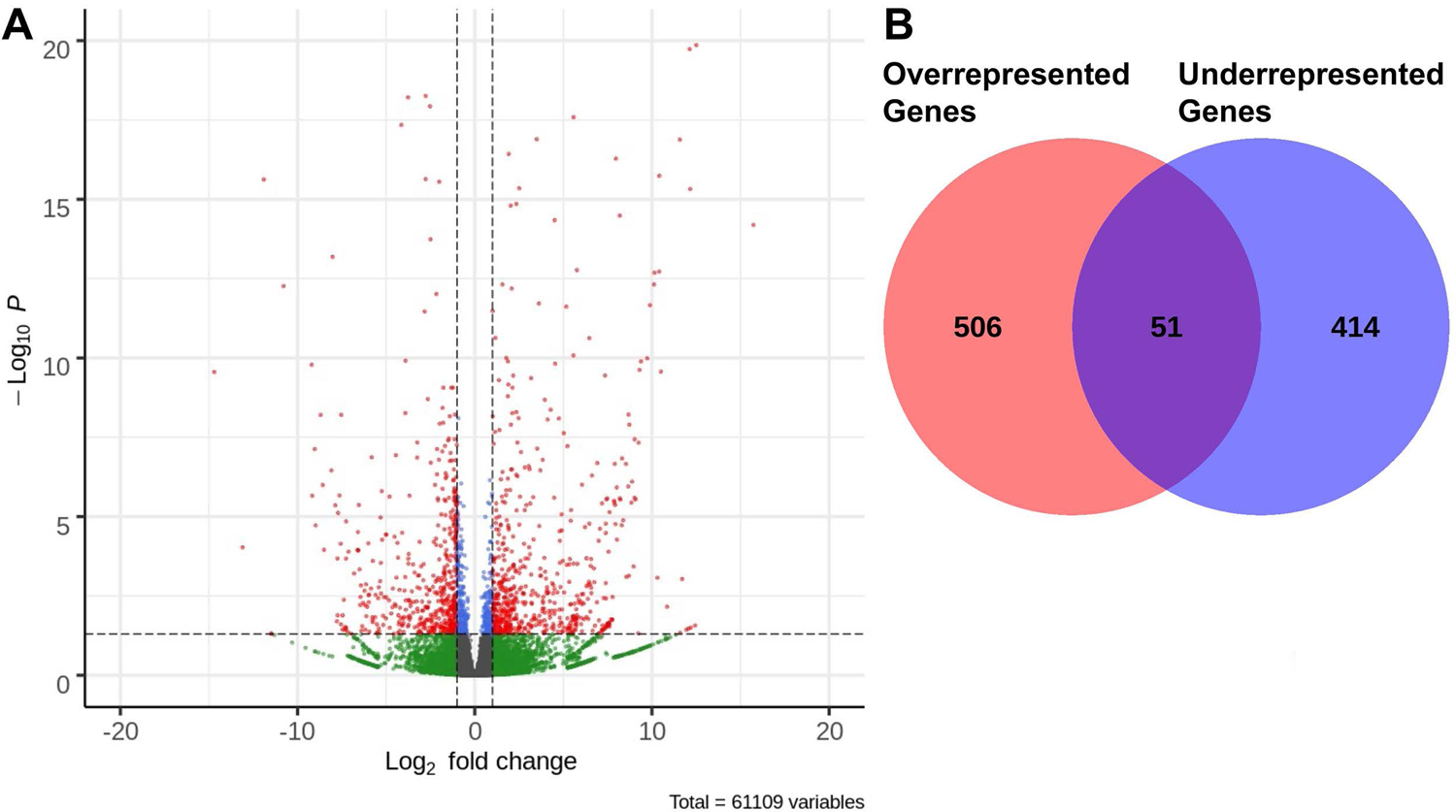
Differential transcript expression in *etr-1(lq61)* using DEseq2. A) A volcano plot of differential transcript expression in *etr-1(lq61)*. Significant cutoff is *q* < 0.05 and log_2_ fold cutoff is log_2_ fold > 1. Transcripts that meet these criteria are indicated by red dots. 630 transcripts were overrepresented, 550 transcripts were underrepresented. B) Venn diagram of genes that had transcripts overrepresented or underrepresented: 506 genes had transcripts that were only overrepresented in *etr-1* muscle cells; 414 genes had transcripts that were only underrepresented; and 51 genes had both over- and underrepresented transcripts.

**Table 1.** Genes with transcripts both overrepresented and underrepresented in etr-1(lq61) muscle.

|  |  |  |
| --- | --- | --- |
| *unc-52* | *pnk-1* | *tag-151* |
| *lev-11* | *sir-2.1* | *mrp-1* |
| *daf-3* | *T10C6.6* | *mbk-2* |
| *rpt-4* | *slo-2* | *T23E7.2* |
| *unc-22* | *spv-1* | *ZK185.1* |
| *dur-1* | *unc-73* | *unc-68* |
| *alp-1* | *lec-3* | *npp-9* |
| *fhod-1* | *cup-5* | *pdi-2* |
| *ntl-2* | *T13H5.1* | *pbrm-1* |
| *ttn-1* | *unc-16* | *deb-1* |
| *yif-1* | *tnt-3* | *add-2* |
| *atn-1* | *slo-1* |  |
| *egl-2* | *F49E2.5* |  |
| *ceh-79* | *unc-36* |  |
| *nsy-7* | *ret-1* |  |
| *avr-14* | *dnj-5* |  |
| *mel-11* | *pqn-80* |  |
| *rme-1* | *tax-6* |  |
| *nhre-61* | *F25E5.8* |  |
| *T07D3.9* | *tep-1* |  |

Gene Ontology enrichment analysis was conducted on four separate groups of these muscle genes with transcripts affected by *etr-1(lq61)* (Supplemental File 5): all genes; genes with transcripts that were only underrepresented; genes transcripts that were only overrepresented; and genes that had transcripts both over- and underrepresented. The six most significant GO terms for each group are shown in Figure 5. Considering all genes, GO terms associated with translation and ribosomal function were apparent, as well as myofilament structure, muscle physiology, reproduction, and embryonic and larval growth (Figure 5A). Genes with transcripts underrepresented in *etr-1(lq61)* were described by GO terms representing translation and ribosomal function (10 of the 18 top GO terms) (Figure 5B). Genes with overrepresented transcripts were described by GO terms representing a broad cross section of cellular function, but translation and the ribosome were not among these (Figure 5C). Genes with both over- and underrepresented transcripts were described by GO terms representing myofilament lattice and muscle physiology, and other cellular functions (Figure 5D). In sum, this GO term analysis suggests that *etr-1(lq61)* influences a broad spectrum of cellular events in muscle, including myofilament lattice and muscle physiology, as well as translation and ribosomal function. Notably, genes involved in translation and ribosomal function are strongly represented among those with transcripts reduced in *etr-1(lq61)*. A similar reduction of expression of genes involved in translation and ribosomal function was described after siRNA knock-down of *CELF1* in chicken cardiomyocytes (Blech-Hermoni *et al*. 2016).

**Figure 5.**
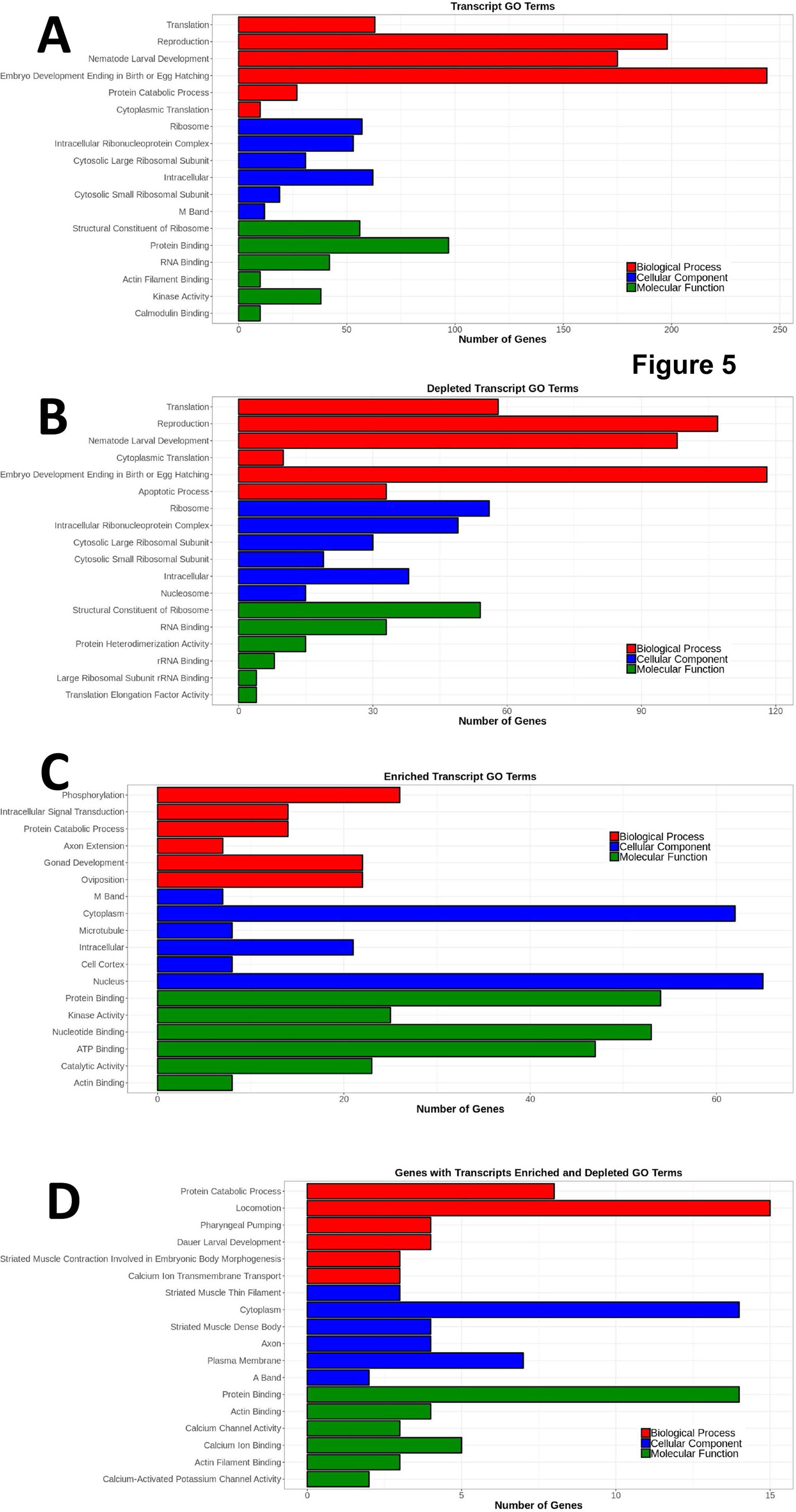
Gene ontology (GO) term analysis of genes with differential transcript representation in *etr-1(lq61)*, as described in Figure 3. The six most significant GO terms (*q* ≤ 0.05) for three categories (biological process, cellular component, and molecular function) are shown. A) All genes with transcripts affected by *etr-1(lq61).* B) Genes with underepresented transcripts. C) Genes with overrepresented transcripts. D) Genes with both underrepresented and overrepresented transcripts. The entire GO term analysis is in Supplemental File 5.

### Genes identified by both differential exon usage and transcript accumulation

*etr-1(lq61)* affected transcript expression of 971 genes, and alternative exon representation of 244 genes. There were 102 genes shared between the 244 alternatively-spliced genes and the 971 genes that had differentially-expressed transcripts (Table 3), a significant association (*p < 0.0001)*. These might represent genes with transcripts for which ETR-1 controls both splicing and transcript accumulation. Alternately, the differential use of exons could influence the transcript accumulation DEseq2 algorithm, leading to under- or overrepresentation of transcripts by alternate exon usage. In any event, identification in both analyses suggests that ETR-1 might have strong effects on the transcripts of these genes.

### Genes affecting AQR and PQR migration

This analysis suggests that ETR-1 regulates multiple aspects of muscle cell function, most notably myofilament lattice, muscle physiology, and translation and ribosomal function. Previous studies indicated that *etr-1(lq61)* had a muscle-derived, non-autonomous effect on migration of AQR and PQR neurons (Ochs *et al*. 2020). Thus, ETR-1 might regulate a secreted signal from the muscles that directs AQR and PQR migration. Genetic analysis suggests this signal could act with or in parallel to Wnt signaling (Ochs *et al*. 2020), which directs AQR and PQR migration (Josephson *et al*. 2016a).

We used feeding RNAi (see Materials and Methods) and mutants to knock down a subset of genes with known roles in cell migration, and genes that had transcripts both overrepresented and underrepresented in *etr-1(lq61)* muscles versus wild-type muscles (Table 1). AQR and PQR position in these animals was scored (see Materials and Methods) (Table 3). In *unc-52, unc-71,* and *skn-1,* between 2% and 5% of AQR and PQR neurons migrated in the wrong direction, as evidenced by AQR in position 5, and PQR in positions 1, 2, and 3. UNC-52 is the basement membrane heparan sulfate proteoglycan Perlecan (Rogalski *et al*. 1993) and will be discussed in more detail below. UNC-71 is an ADAM metalloprotease that has been shown to act in anterior Q descendant migration (Huang *et al*. 2003; Masuda *et al*. 2012). SKN-1 is an ortholog of the human NFE2L1 transcriptional regulator that controls a wide variety of developmental events including muscle differentiation (Gomes *et al*. 2001; Broitman-Maduro *et al*. 2005). Some genes displayed defects in the ability of AQR and PQR to migrate (>10%), but not directional defects, as evidenced by AQR in positions 2, 3, and 4 and PQR in position 4. Many genes displayed few (<10%) or no defects in the ability of AQR and PQR to migrate. Thus, some genes with transcripts that are regulated by ETR-1 in muscle had instructional roles in directing AQR and PQR migration, and some had permissive roles in the ability of AQR and PQR to migrate.

### The heparan sulfate proteoglycan UNC-52 affects AQR and PQR migration

*unc-52* encodes the basement membrane heparan sulfate proteoglycan Perlecan and is involved in myofilament lattice attachment to the basement membrane (Rogalski *et al*. 1993). *unc-52* is extensively alternatively spliced (Rogalski *et al*. 1995; Rogalski *et al*. 2001), including in the epidermis by CCAR-1 (Fu *et al*. 2018) and MEC-8 (Lundquist *et al*. 1996; Spike *et al*. 2002) with consequences on hemidesmosome formation, muscle attachment and mechanosensory neuron function.

We found that *etr-1(lq61)* affected *unc-52* transcript expression in muscle cells. *unc-52* was identified in both exon representation by DEXseq and transcript accumulation by DEseq2 (Table 2). In *etr-1(lq61)* muscle cells, some 5’ exons were significantly overrepresented and 3’ exons significantly underrepresented compared to *wild-type* muscle cells. The far 5’ exons (see Figure 6A) predicted for *unc-52* were not highly expressed in muscle in either background. We visualized *unc-52* splice junctions using the Sashimi plot function in the Integrated Genome Viewer (see Materials and Methods) (Figure 6B). In wild-type animals, 3’ exons were well-represented (Figure 6B). In *etr-1(lq61)*, 3’ exons were significantly underrepresented and 5’ exons significantly overrepresented. These data suggest that ETR-1 proteins that include exon 8 are required to produce the long isoforms of *unc-52* containing the 3’ exons in muscles.

**Figure 6.**
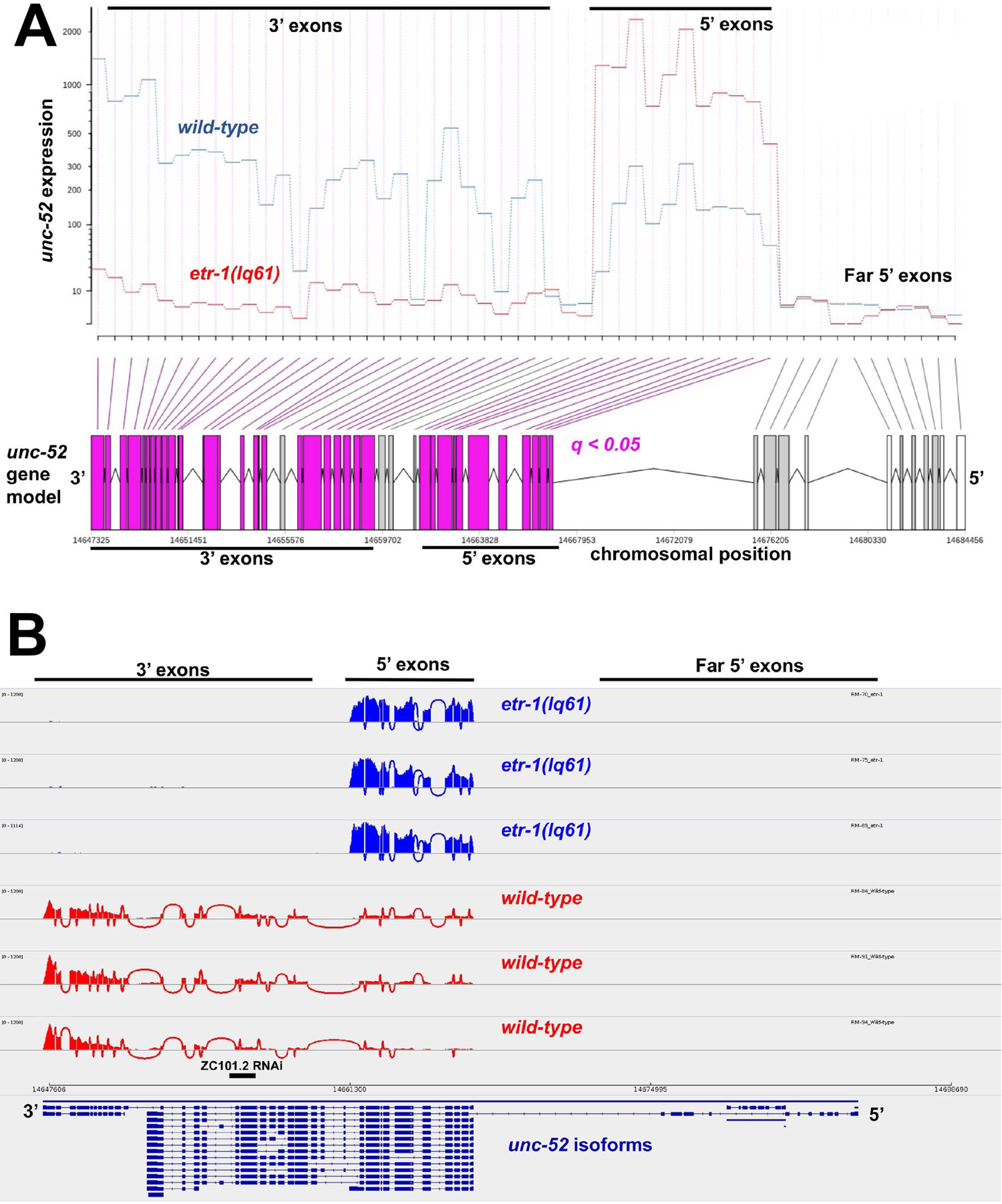
*etr-1(lq61)* affects *unc*-*52* exon usage. A) DEXSeq output comparing *etr-1(lq61)* muscle to wild-type muscle as described in Figure 2. B) An IGV-Sashimi plot comparing *unc-52* exon usage in *etr-1(lq61)* muscle cells and wild-type muscles. The blue peaks represent *etr-1(lq61)* splice junctions in muscle cells in three independent biological replicates, and the red peaks represent splice junctions in wild-type muscles in three independent biological replicates.

**Table 2.** Genes with both exon representation and transcript accumulation affected by etr-1(lq61).

|  |  |  |  |  |
| --- | --- | --- | --- | --- |
| *atn-1* | *mrp-1* | *unc-73* | *cpna-2* | ZK370.4 |
| *avr-14* | *nhr-61* | *unc-87* | *wht-1* | *nsy-7* |
| *ccr-4* | *ntl-2* | *unc-89* | *fln-2* |  |
| *ced-1* | *num-1* | *unc-96* | *crh-2* |  |
| *ctn-1* | *ketn-1* | *ver-1* | C27H5.2 |  |
| *deb-1* | *ret-1* | *zmp-1* | *pxl-1* |  |
| *dur-1* | *rme-1* | *alx-1* | *tag-250* |  |
| *alp-1* | *rpl-19* | C06B3.6 | *acs-17* |  |
| *egl-10* | *sli-1* | *ceh-79* | *fhod-1* |  |
| *etr-1* | *slo-1* | *tbc-19* | C56G2.9 |  |
| *gcy-31* | *srp-7* | *pde-3* | F13C5.5 |  |
| *gld-2* | *lim-9* | *shw-1* | F20A1.6 |  |
| *gon-1* | *ttn-1* | F23B12.7 | *dip-2* |  |
| *hsp-4* | *dgk-4* | *pole-1* | F44E7.4 |  |
| *kin-1* | *cpna-1* | *mlcd-1* | F46F11.1 |  |
| *lea-1* | *tns-1* | F40F8.5 | F48E3.8 |  |
| *lec-2* | *tax-6* | F43G9.12 | K09F6.10 |  |
| *lec-3* | *tmd-2* | F49E2.5 | *trx-3* |  |
| *lev-11* | *tnt-3* | *yif-1* | M01H9.3 |  |
| *lin-45* | *tsp-17* | *magi-1* | R11G1.6 |  |
| *lip-1* | *unc-32* | *nep-21* | T23E7.2 |  |
| *lsm-5* | *unc-36* | T28F4.5 | *zig-11* |  |
| *mai-1* | *unc-52* | *grdn-1* | *fln-1* |  |
| *mel-11* | *unc-64* | Y105E8A.25 | *him-19* |  |
| *mlp-1* | *unc-68* | *nit-1* | ZC449.5 |  |

As described above, *unc-52(RNAi)* resulted in AQR and PQR directional migration defects (Table 1). For RNAi, we utilized the ZC101.2 Source Bioscience II-9A20 clone, which is located located in the 3’ region of *unc-52*, the region that is underrepresented in in *etr-1(lq61)* muscles (Figure 6B). *unc-52(RNAi)* animals displayed the <u>p</u>aralyzed, <u>a</u>rrested at two-fold stage (Pat) phenotype similar to strong loss-of-function alleles of *unc-52* (Figure 7A and B) (Williams and Waterston 1994). Notably, *etr-1(lq61)* animals, which do not express the 3’ exons from this *unc-52* region in muscle, do not show the Pat phenotype. The viability of *etr-1(lq61)* mutants could be due to a muscle-specific effect of ETR-1 on *unc-52*, with *unc-52* long isoforms with 3’ exons expressed and functional in other tissues (*e.g.* hypodermis) in *etr-1(lq61)* mutants.

**Figure 7.**
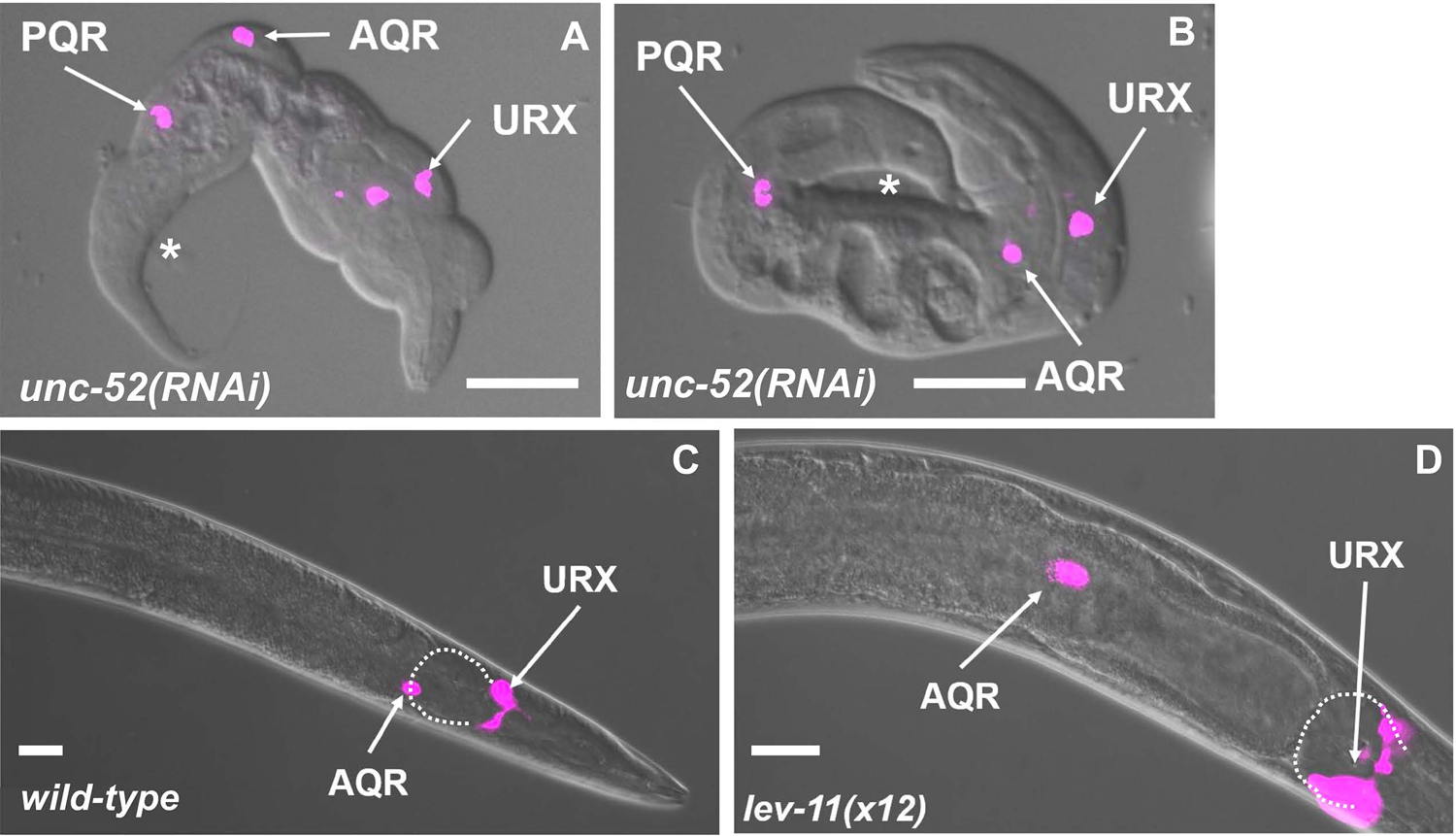
AQR and PQR defects in RNAi knockdown of *unc-52* and *lev-11*. Micrographs of merged DIC and fluorescence images are shown, with *gcy-32::cfp* expression in AQR and PQR (magenta). Two representative examples of paralyzed arrested at two-fold stage (Pat) *unc-52(RNAi)* animals are shown. The anus is indicated by an asterisk. A) An *unc-52(RNAi)* animal displayed misplaced AQR and PQR. B) An *unc-52(RNAi)* animal displayed a misplaced PQR, with AQR in the normal position. C) The wild-type position of AQR just posterior to the posterior pharyngeal bulb (outlined with a dashed white line). B) AQR position in a *lev-11(x12)* mutant. AQR was displaced posteriorly approximately 50μm from the posterior pharyngeal bulb. Scale bars represent 10μm.

Despite embryonic lethality and the Pat phenotype, AQR and PQR were visible in *unc-52(RNAi)* arrested Pat animals. AQR and PQR displayed defects in the ability of AQR and PQR to migrate (Figure 7A and B) as well as defects in direction of migration (Table 3). In sum, ETR-1 proteins that contain exon 8 are required for the accumulation of long isoform transcripts of *unc-52* containing 3’ exons in muscles. Targeting the 3’ region of *unc-52* with RNAi resulted in AQR and PQR defects, suggesting a role in AQR and PQR migration.

**Table 3.**
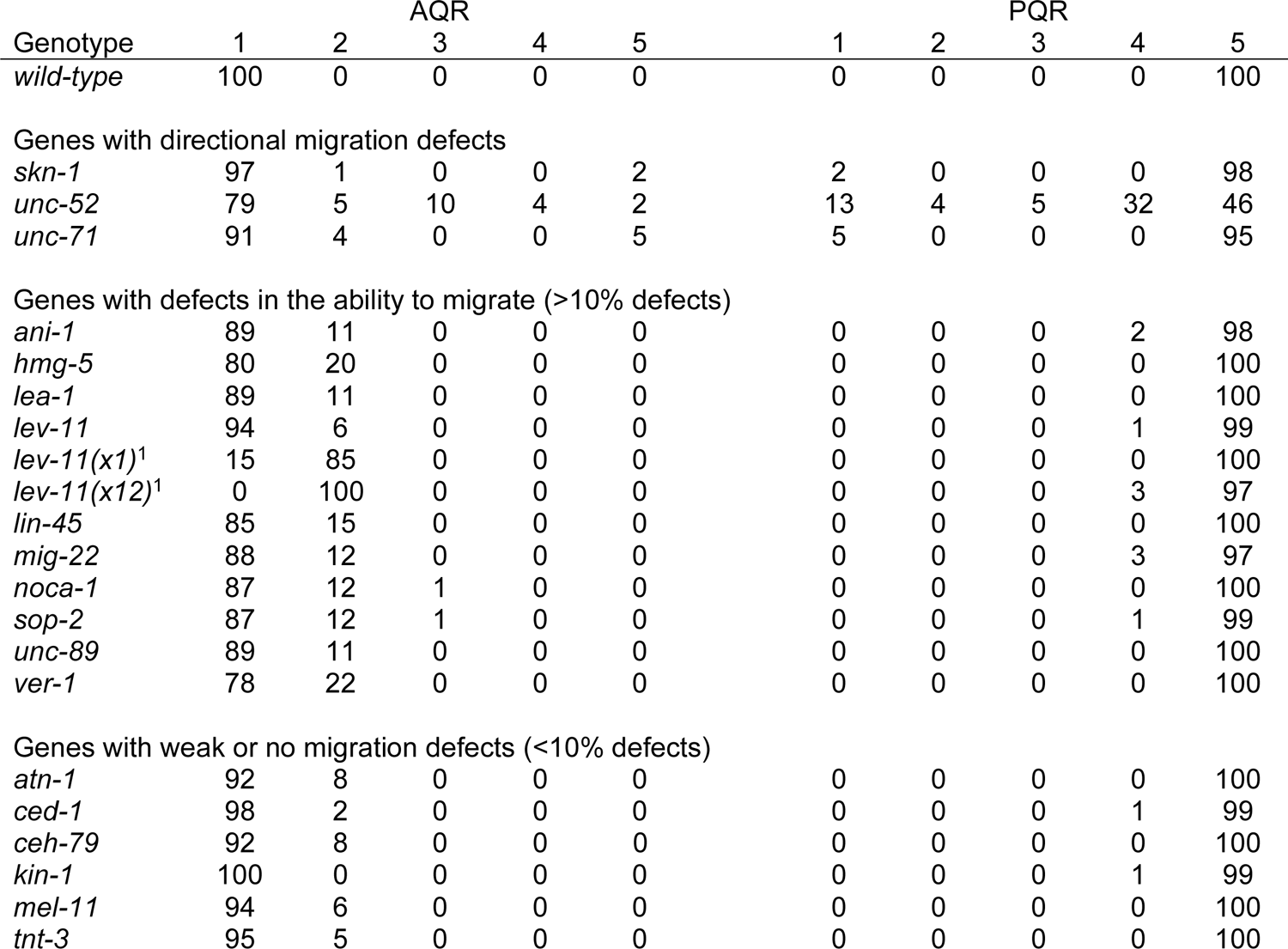
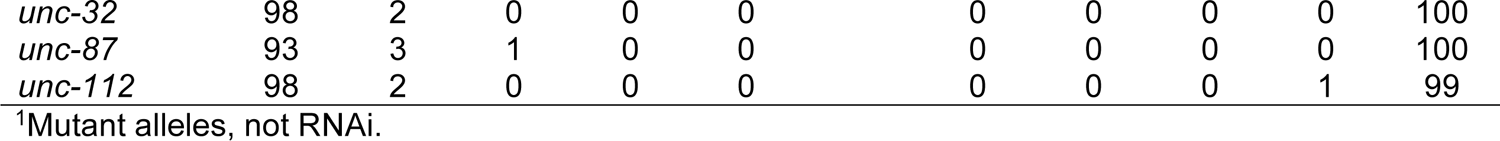
Candidate gene AQR and PQR migration defects using RNAi unless otherwise noted.

| Genotype | AQR |  |  |  |  | PQR |  |  |  |  |
| --- | --- | --- | --- | --- | --- | --- | --- | --- | --- | --- |
|  | 1 | 2 | 3 | 4 | 5 | 1 | 2 | 3 | 4 | 5 |
| *wild-type* | 100 | 0 | 0 | 0 | 0 | 0 | 0 | 0 | 0 | 100 |
| Genes with directional migration defects |  |  |  |  |  |  |  |  |  |  |
| *skn-1* | 97 | 1 | 0 | 0 | 2 | 2 | 0 | 0 | 0 | 98 |
| *unc-52* | 79 | 5 | 10 | 4 | 2 | 13 | 4 | 5 | 32 | 46 |
| *unc-71* | 91 | 4 | 0 | 0 | 5 | 5 | 0 | 0 | 0 | 95 |
| Genes with defects in the ability to migrate (>10% defects) |  |  |  |  |  |  |  |  |  |  |
| *ani-1* | 89 | 11 | 0 | 0 | 0 | 0 | 0 | 0 | 2 | 98 |
| *hmg-5* | 80 | 20 | 0 | 0 | 0 | 0 | 0 | 0 | 0 | 100 |
| *lea-1* | 89 | 11 | 0 | 0 | 0 | 0 | 0 | 0 | 0 | 100 |
| *lev-11* | 94 | 6 | 0 | 0 | 0 | 0 | 0 | 0 | 1 | 99 |
| *lev-11(x1)<sup>1</sup>* | 15 | 85 | 0 | 0 | 0 | 0 | 0 | 0 | 0 | 100 |
| *lev-11(x12)<sup>1</sup>* | 0 | 100 | 0 | 0 | 0 | 0 | 0 | 0 | 3 | 97 |
| *lin-45* | 85 | 15 | 0 | 0 | 0 | 0 | 0 | 0 | 0 | 100 |
| *mig-22* | 88 | 12 | 0 | 0 | 0 | 0 | 0 | 0 | 3 | 97 |
| *noca-1* | 87 | 12 | 1 | 0 | 0 | 0 | 0 | 0 | 0 | 100 |
| *sop-2* | 87 | 12 | 1 | 0 | 0 | 0 | 0 | 0 | 1 | 99 |
| *unc-89* | 89 | 11 | 0 | 0 | 0 | 0 | 0 | 0 | 0 | 100 |
| *ver-1* | 78 | 22 | 0 | 0 | 0 | 0 | 0 | 0 | 0 | 100 |
| Genes with weak or no migration defects (<10% defects) |  |  |  |  |  |  |  |  |  |  |
| *atn-1* | 92 | 8 | 0 | 0 | 0 | 0 | 0 | 0 | 0 | 100 |
| *ced-1* | 98 | 2 | 0 | 0 | 0 | 0 | 0 | 0 | 1 | 99 |
| *ceh-79* | 92 | 8 | 0 | 0 | 0 | 0 | 0 | 0 | 0 | 100 |
| *kin-1* | 100 | 0 | 0 | 0 | 0 | 0 | 0 | 0 | 1 | 99 |
| *mel-11* | 94 | 6 | 0 | 0 | 0 | 0 | 0 | 0 | 0 | 100 |
| *tnt-3* | 95 | 5 | 0 | 0 | 0 | 0 | 0 | 0 | 0 | 100 |
| *unc-32* | 98 | 2 | 0 | 0 | 0 | 0 | 0 | 0 | 0 | 100 |
| *unc-87* | 93 | 3 | 1 | 0 | 0 | 0 | 0 | 0 | 0 | 100 |
| *unc-112* | 98 | 2 | 0 | 0 | 0 | 0 | 0 | 0 | 1 | 99 |

<sup>1</sup>Mutant alleles, not RNAi.

### RNAi knockdown of *lev-11* results in AQR and PQR migration defects

*lev-11* was also identified by both exon representation and transcript accumulation in *etr-1(lq61)* (Table 2). *lev-11* encodes a tropomyosin (Kagawa *et al*. 1997), which is a known target of vertebrate CELF1 (Dasgupta and Ladd 2012). *lev-11* encodes multiple isoforms regulated in a tissue-specific manner (Anyanful *et al*. 2001; Watabe *et al*. 2018).

*lev-11* exons 8 and 15 were significantly overrepresented in *etr-1(lq61)* muscle compared to wild-type muscle (Figure 8A). IGV-Sashimi splice junction analysis revealed that *lev-11* exons 8 and 15 were included in *etr-1(lq61)* muscle and largely excluded in wild-type muscle (Figure 8B). These results suggest that ETR-1 proteins with exon 8 controlled alternative splicing of *lev-11* in muscle cells, most notably removing exons 8 and 15 from *lev-11* transcripts.

**Figure 8.**
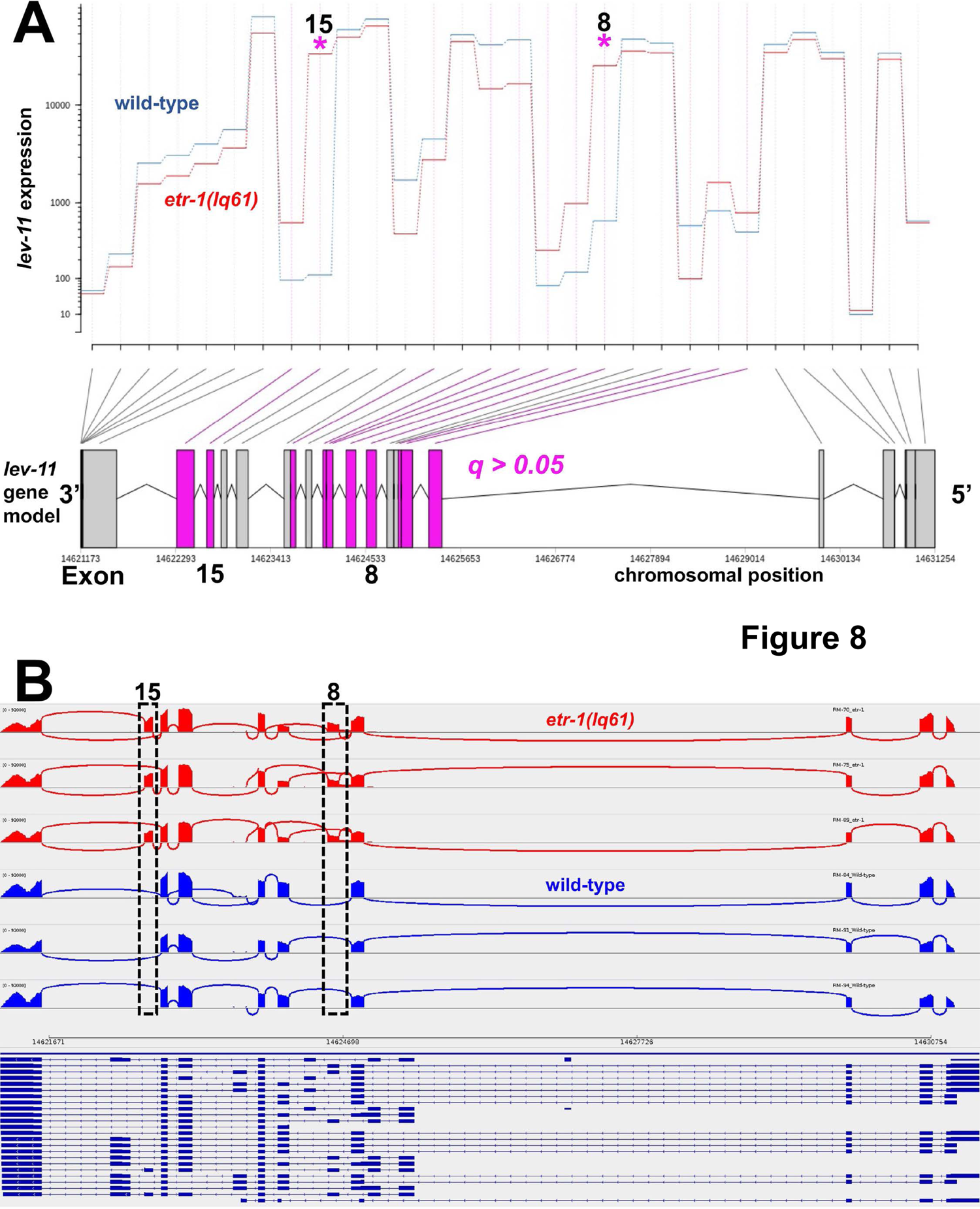
*etr-1(lq61)* affects *lev-11* exon usage. A) DEXSeq output for *lev-11* comparing exon usage levels between *etr-1(lq61)* muscle and wild-type muscle, as described in Figure 2. B) IGV-Sashimi plot of *lev-11* splice junctions in wild-type and *etr-1(lq61*), as described in Figure 6.

*lev-11* RNAi resulted in AQR not migrating the full distance anteriorly, a phenotype also observed in *lev-11* mutants (Table 3 and Figure 7C and D). The region at which AQR stopped migrating in both *lev-11* RNAi and *lev-11* mutants was highly stereotyped, in position 4 just posterior to the normal position. These results suggest that *lev-11* might affect a specific developmental sign post rather than the general ability of AQR to migrate. In sum, our functional analysis indicates that genes with transcripts regulated by ETR-1 in muscles include those that control AQR and PQR neuron migration.

## Discussion

### Identification of ETR-1/CELF target genes via alternative exon usage and differential transcript abundance

CELF-family proteins regulate mRNA processing, including alternative splicing and transcript stability (reviewed in (Dasgupta and Ladd 2012)). The *etr-1(lq61)* mutation presented a unique opportunity to determine CELF target genes in a specific tissue, body wall muscles of *C. elegans*. First, *lq61* corresponds to a premature stop codon in an alternatively spliced exon affecting only a subset of *etr-1* transcripts, and thus did not cause embryonic lethality (Ochs *et al*. 2020). Because *etr-1(lq61)* animals are viable, it was possible to isolate body wall muscle cells and their transcripts at a specific timepoint in L1 animals when Q neuroblasts are migrating. Second, *etr-1(lq61)* caused defects in AQR and PQR neuron migration in a cell-non-autonomous manner (Ochs *et al*. 2020). Thus, ETR-1 targets might be involved in producing a guidance signal for neurons from body wall muscle cells. Such an interaction might be conserved in vertebrate CELF function.

Two algorithms were used to compare differences in the transcriptomes of wild-type and *etr-1(lq61)* animals. DEXseq was used to assay alternative exon representation, and DEseq2 was used to assay differential transcript representation and splice junction usage. This analysis identified 244 genes with significant differential exon representation, and 1180 transcripts, corresponding to 971 genes, that were differentially represented. 102 genes were identified in both analyses, suggesting that these are not discrete categories and that exon abundance can affect predicted transcript abundance and *vice versa*. Our results suggest that a gene-by-gene approach using output from DEXseq, DEseq2, and IGV-Sashimi gives a clear picture of the effects on transcript structure and abundance.

It is likely that other ETR-1 targets were not identified in this analysis, as only a subset of *etr-1* isoforms containing exon 8 are affected by *etr-1(lq61)*. Exon 8 encodes a polyglutamine-rich region (Ochs *et al*. 2020), which is thought to mediate interaction of CELF proteins with other splicing/RNA processing factors thus potentially altering the efficacy and specificity of the splicing reaction (SINGH *et al*. 2011). The genes identified in this analysis could be direct targets of ETR-1, or indirect targets through the effects of ETR-1 on other RNA processing genes.

### ETR-1 target genes include those involved in myofilament lattice assembly and function, muscle cell physiology, and translation

GO term analysis of ETR-1 target genes identified broad effects on the muscle cell transcriptome. The most significant GO terms described genes involved in myofilament lattice structure, assembly, and attachment were identified, along with genes involved in muscle cell physiology. This is consistent with the Pat phenotype of *etr-1* RNAi (Milne and Hodgkin 1999) and with the known effects of CELF family members in muscle cell physiology. GO terms involving germ cell and reproduction were also defined, consistent with a known role of *etr-1* in germ cell corpse apoptosis (Boateng *et al*. 2017). These genes might have conserved functions in the germline and in muscle.

Strikingly, the most significantly enriched GO terms describing genes with underrepresented transcripts in *etr-1(lq61)* involved translation and ribosomal function. Translation and ribosome GO terms were largely absent from the most significant GO terms describing genes with overrepresented transcripts and alternative exon usage, suggesting translation and ribosome GO terms were specifically associated with genes that require ETR-1 for transcript accumulation. This also suggests a general impairment of translation in *etr-1(lq61)* muscles. siRNA knock-down of *CELF1* in chicken cardiomyocytes also led to decreased expression of coding sequences with GO terms associated with translation and ribosomal function (Blech-Hermoni *et al*. 2016). These parallel findings point to a potentially ancient evolutionary origin of CELF-family targets. Indeed, known targets of CELF molecules include tropomyosin, actinin, and troponin (Suzuki *et al*. 2002; Dasgupta and Ladd 2012). Our analysis identified *lev-11/tropomyosin, atn-1/actinin,* and *tnt-3/troponin* as ETR-1 targets with differential alternative splicing events. Depletion of *etr-1* results in increased apoptotic bodies in the gonad, dependent upon the CED-1 cell corpse scavenger receptor (Boateng and Allen 2018). Our analysis in muscles revealed a significant increase in the *ced-1* transcript in *etr-1(lq61)* (log_2_ fold change 1.58, *q* = 0.002), suggesting that the interaction with CED-1 might be conserved in muscles.

Other known CELF1 targets were not identified in our analysis. For example, *MTMR1* and *MTMR3*, encoding myotubularin-related proteins, are alternatively spliced by CELF1 during mouse heart development (Ladd *et al*. 2005; Kalsotra *et al*. 2008; Santoro *et al*. 2010). However, we did not detect significant differences in exon or transcript abundance of the related *C. elegans* genes, *mtm-1* and *mtm-3*. Possibly, *mtm-1* and *mtm-3* are not regulated by ETR-1 isoforms that contain exon 8. Alternately, ETR-1 might not regulate *mtm-1* and *mtm-3* in body wall muscle in *C. elegans*, as these genes were identified as CELF1 targets in cardiomyocytes (Ladd *et al*. 2005; Kalsotra *et al*. 2008).

### Targets shared between ETR-1 and UNC-75

The CELF family of proteins contains six members in mouse and humans (Dasgupta and Ladd 2012). In *C. elegans* there are two members, ETR-1, similar to CELF1-2, and UNC-75, similar to CELF3-6. *unc-75* function is predominantly restricted to the nervous system (Loria *et al*. 2003; Kuroyanagi *et al*. 2013a; Kuroyanagi *et al*. 2013b; Norris *et al*. 2014; Chen *et al*. 2016). Targets of UNC-75 in the nervous system include *unc-32*, which encodes for the α subunit of the V_0_ complex of vacuolar-type H^+^-ATPases (Kuroyanagi *et al*. 2013a), and *unc-64/syntaxin* (Norris *et al*. 2014; Chen *et al*. 2016). Both *unc-32* and *unc-64* were identified as ETR-1 targets in body wall muscle cells in our analysis. This finding suggests that ETR-1 and UNC-75 might regulate common targets in a tissue-specific manner (i.e. ETR-1 in muscle, and UNC-75 in neurons).

### ETR-1 targets control AQR and PQR neuronal migration

Previous studies showed that ETR-1 acts cell-non-autonomously in muscle to control AQR and PQR neuronal migration (Ochs *et al*. 2020). ETR-1 might be involved in the generation of a signal from body wall muscles that controls AQR and PQR migration. Body wall muscle is known source of cues that control Q neuroblast lineage migration. SPON-1/F-spondin from posterior body wall muscles is required for robust AQR and PQR migration (Josephson *et al*. 2016b). Furthermore, the NFM-1/NF-2 molecule acts non-autonomously, possibly in muscles, to control Q neuroblast protrusion and migration (Josephson *et al*. 2017).

ETR-1 targets analyzed with the strongest effects on AQR and PQR migration include *skn-1, unc-71, unc-52*, and *lev-11*. The ADAM metalloprotease UNC-71 was previously implicated in anterior QR migrations (Huang *et al*. 2003; Masuda *et al*. 2012), and our results suggest that UNC-71 also controls directional PQR migration. SKN-1, UNC-52, and LEV-11 have not been previously implicated in Q lineage migration. The NFE2L1 transcriptional regulator SKN-1 is a known regulator of muscle differentiation (Gomes *et al*. 2001; Broitman-Maduro *et al*. 2005). While RNAi knockdown of these genes in all tissues by feeding RNAi caused AQR and PQR migration defects, further studies will be required to show that ETR-1 regulation of these genes in muscle cells is relevant to AQR and PQR migration (i.e. these genes might act in other cells besides muscles to control AQR and PQR migration).

### *unc-52*/Perlecan is a target of *etr-1* and is required for AQR and PQR migration

*unc-52* encodes the basement membrane heparan sulfate proteoglycan Perlecan and is expressed in many different cell types, including the muscle cells (Rogalski *et al*. 1993; Mullen *et al*. 1999; Spike *et al*. 2002). *unc-52* has been shown to control the migrations of the distal tip cells (Merz *et al*. 2003). The heparan sulfate epimerase HSE-5 has been shown to control Q neuroblast migration (Sundararajan *et al*. 2015; Wang *et al*. 2015), but loss of no single HSPG or in double mutant combination has been shown to affect Q migrations, including hypomorphic *unc-52* alleles (Sundararajan *et al*. 2015).

*unc-52* was identified as a target of ETR-1 in muscles, with the 3’ exons of *unc-52* underrepresented in *etr-1(lq61)*. This finding suggests that ETR-1 with exon 8 is required for the accumulation of the long *unc-52* isoforms with the 3’ exons. Depletion of *unc-52* by RNAi resulted in embryonic lethality and severe AQR and PQR migration defects, including directional defects similar to *hse-5*. This finding suggests that UNC-52 might be the HSPG through which HSE-5 is controlling Q migration. Both *unc-52* and *hse-5* mutants display defects in AQR and PQR direction of migration, suggesting that UNC-52 is not merely a substrate for cell migration but rather provides directional information.

Our analysis showed that *unc-52* isoforms containing the 3’ exons were strongly underrepresented in *etr-11(lq61)* muscle, with the shorter ZC101.2b.1 transcript the predominant *unc-52* mRNA. The *unc-52* RNAi clone we used is located in these 3’ exons, suggesting that the long isoforms with the 3’ exons are important for AQR and PQR migration. The 3’ exons mainly encode for Immunoglobulin family domains, some of which are alternatively spliced in the epidermis by other factors, MEC-8 and CCAR-1 (Lundquist *et al*. 1996; Spike *et al*. 2002; Fu *et al*. 2018). RNAi of *unc-52* resulted in embryonic lethality, yet *etr-1(lq61)*, which nearly eliminates transcripts with the 3’ exons, is viable. This is likely due to *unc-52* function in other tissues not regulated by *etr-1* (i.e. epidermis). Indeed, elimination of transcripts with the 3’ exons in the epidermis results in embryonic lethality in *mec-8; unc-52(hypomorphic)* double mutants (Spike *et al*. 2002). This suggests that *unc-*52 long isoforms in muscle are not required for embryonic viability but are required for AQR and PQR migration.

### *lev-11* is alternatively spliced by *etr-1* and is required for AQR and PQR migration

We showed that ETR-1 exon 8 isoforms are required for alternative splicing of *lev-11* transcripts in muscle by the removal of exons 8 and 15 (Figure 8B). It was shown previously that CELF proteins regulate the alternative splicing of tropomyosin in chicken muscle development (Sureau *et al*. 2011). Isoforms of *lev-11* are expressed in a tissue-specific manner (Watabe *et al*. 2018). *lev-11* isoforms without exons 8 and 15 are expressed robustly in wild-type body wall muscle cells (Watabe *et al*. 2018) (Figure 8B), whereas those containing *lev-11* exons 8 and 15 are expressed in pharyngeal muscle and the excretory cell (Watabe *et al*. 2018). Our results suggest that ETR-1 with exon 8 is required to exclude exons 8 and 15 from *lev-11* transcripts in body wall muscle cells. Possibly, *lev-11* isoforms lacking exons 8 and 15 are optimized for wall body muscle structure or function.

Knockdown of *lev-11* by either RNAi or in *lev-11* mutants resulted in AQR migration defects, with most AQR cells stopping approximately 50μm posterior to its normal position near the posterior pharyngeal bulb. This characteristic stopping point suggests that *lev-11* affects a developmental guidepost rather than generally affecting the ability of AQR to migrate, which results in stoppages of AQR along anterior body in positions 2, 3, and 4 (*e.g. epi-1/laminin* mutants) (Lang and Lundquist 2021). Two-hybrid studies suggest that LEV-11 physically interacts with MIG-14/Wntless (Koorman *et al*. 2016), which is involved involved in Wnt secretion (Banziger *et al*. 2006; Pan *et al*. 2008; Sawa and Korswagen 2013). Mutation of *mig-14* results in Q neuroblast migration defects (Maloof *et al*. 1999; Ch’ng *et al*. 2003; Harterink *et al*. 2011), including AQR and PQR directional defects (Josephson *et al*. 2016a). *etr-1* interacts genetically with *Wnt* mutations to control AQR and PQR migration, either directly or through a parallel pathway (Ochs *et al*. 2020). Expression of three *Wnt* genes is significantly increased in body wall muscle (Supplemental File 1): *egl-20* (log2-fold change = 2.96), *cwn-1* (log2-fold change = 2.74), and *cwn-2* (log2-fold change = 1.02), but their exon use and transcript levels are not affected by *etr-1(lq61)*. Intriguingly, *cwn-1* mutants display posteriorly-displaced AQR similar to but weaker than *lev-11* mutants (Josephson *et al*. 2016a). It is plausible that *etr-1* regulates the splicing of *lev-11* in muscle to control an interaction with MIG-14. Disrupting this interaction might result in improper production of a Wnt signal, causing AQR and PQR migration defects. Further studies will be aimed at exploring this mechanism.

### Linking the effects of *etr-1(lq61)* on the muscle cell transcriptome to mutant phenotypes

Here we define transcripts in body wall muscle cells that are affected by mutation of the CELF1 family member *etr-1*. The *etr-1(lq61)* mutation selectively disrupts isoforms that include exon 8, leaving other ETR-1 isoforms intact. Thus, this set of body muscle transcripts include only those affected by ETR-1 with exon 8. *etr-1(lq61)* affected splicing and accumulation of transcripts of genes involved in a broad spectrum of muscle cell function including myofilament structure and attachment and physiology. Strikingly, genes involved in translation and ribosomal function were abundant among genes with transcripts underrepresented in *etr-1(lq61)*. siRNA knock-down of *CELF1* in chicken cardiomyocytes also led to a decrease in genes with GO terms associated with translation and ribosomal function (Blech-Hermoni *et al*. 2016), suggesting a deep evolutionary conservation of CELF1 target genes. Indeed, *lev-11/tropomyosin, atn-1/actinin,* and *tnt-3/troponin* were also conserved targets in *C.* elegans and vertebrates (Suzuki *et al*. 2002; Dasgupta and Ladd 2012).

How do these broad effects on the muscle transcriptome result in *etr-1(lq61)* mutant phenotypes, including AQR and PQR migration? We demonstrate that genes identified here include those that control AQR and PQR migration, such as *unc-52* and *lev-11*. What is less clear is how the *etr-1-*specific effects on these transcripts in muscle cells contributes to AQR and PQR migration defects. It seems unlikely that the AQR and PQR defects in *etr-1(lq61)* result from disruption of one or a small handful of genes.

Rather, more likely is that subtle effects on multiple genes controlling AQR and PQR migration contribute to the phenotype, such as *unc-52, lev-11*, and others. Future studies will be aimed at addressing this question.

## Materials and Methods

### Strains and Genetics

*C. elegans* strains were was cultured using standard methods at 20 °C. Wild-type strain was N2 Bristol. Alleles used were: LGII: *etr-1(lq61)*, *lqIs244[Pgcy-32::cfp], lex-11(x1 and x12).* LG V: *lqIs58[[Pgcy-32::cfp].* LG unknown: *ccIs4521*

### Larval disruption for cell sorting

For body wall muscle cell sorting, the *myo-3::gfp* transgene *ccIs4521* was used, in *wild-type* and *etr-1(lq61)* backgrounds. *C. elegans* strains were grown on twenty x 150 mm 8P nutrient agar plates (for 1 liter: 20 g bactopeptone, 3 g NaCl, 25 g agar) with *E. coli* strain NA22, which produces a thick bacterial lawn. Early L1 animals were synchronized by first collecting embryos using bleach (hypochlorite) treatment of gravid adult hermaphrodites, and allowing the embryos to hatch overnight into sterile M9 medium at 20°C. This produced approximately 3 million starved, synchronized early L1 larvae. The Q neuroblasts of starved L1s were variably arrested in their early development, from initial migration to the first division after migration (data not shown). Preparations of dissociated larval cells were generated as previously described using SDS-DTT and Pronase treatment coupled with mechanical disruption using a pipet (Spencer *et al*. 2014; Taylor *et al*. 2020).

### FACS analysis

Fluorescence activated cell sorting of *myo-3::GFP-*expressing body wall muscle cells was performed as previously described using a BD FACSAria with a 70 micron diameter nozzle (Spencer *et al*. 2014; Taylor *et al*. 2020). DAPI was added to mark dead or damaged cells which were excluded from each sample of viable myo-3::GFP labeled muscle cells. Profiles of GFP strains were compared to a non-transgene-bearing N2 standard to exclude auto-fluorescent cells. Cells were sorted into Trizol LS. RNA was extracted from the aqueous phase using 5PRIME phase lock gel heavy tubes and purified on a spin column (Zymo, R1013). 30,000-50,000 FACS-sorted cells from each experiment, yielded 5-100 ng total RNA which was used in RNA seq library preparation. Three independent replicate samples were obtained for both wild-type and *etr-1(lq61)* mutant animals.

### mRNA Library Preparation and Sequencing Methods

For an all-cells control group, total RNA was isolated from an aliquot of ∼25,000 L1 larvae set aside before cell dissociation and FACS for each one of the samples. L1 larvae were quickly frozen in liquid nitrogen in a mortar and pestle and ground to powder. Trizol LS was added to the powdered animals for RNA extraction. Muscle FACS and whole L1 RNA extraction resulted in twelve RNA samples representing three biological replicates of N2 and *etr-1(lq61)* sorted muscles and three biological replicates of N2 and *etr-1(lq61)* whole L1 larvae all-cell controls. Quality of total RNA samples was ensured using an Agilent TapeStation, and quantity was determined using Qubit fluorimetry. Sequencing libraries were constructed with 5-100ng of total RNA using the NEBNext Ultra II Directional RNA Library Kit for Illumina. The sequencing library construction process aimed for 300-bp inserts and included mRNA purification with poly-A beads, fragmentation, strand specific cDNA synthesis, end repair, 3’ end adenylation, adapter ligation, and PCR amplification. The constructed sequencing libraries were validated and quantified with Qubit and TapeStation assays. Each library indexed and sequenced in multiplex on the Illumina NextSeq550 system, generating paired-end, 150-base sequence reads from the libraries. Between 33 million and 39 million reads were generated for each of the twelve samples. Base calling was carried out by the Nextseq550 instrument Real Time Analysis (RTA) software. The base call files (bcl files) were demultiplexed and converted to compressed FASTQ files by bcl2fastq2.

### Data Availability Statement

All data are represented in this manuscript and all strains and reagents are available upon request. FASTQ files for this project are available in the Sequence Read Archive (SRA) under project number PRJNA733501. Computational code can be found in Supplemental Materials.

### RNA-Seq Analysis

The *C. elegans* reference genome was downloaded from the following URL: //hgdownload.cse.ucsc.edu/goldenPath/ce11/bigZips/chromFa.tar.gz. Quality control of reads was performed using FastQC (version 0.11.5) and fastp (options: -- detect_adapter_for_pe –length_required 75 –trim_front1 10 –trim_front2 10 – cut_mean_quality 25 –cut_window_quality 5-cut_tail, version 0.19.8) (Chen *et al*. 2018). Splicesite and exon models were built using the *C. elegans* release 11 GTF file and HISAT2 with default settings (hisat2_extract_splice_sites.py and hisat2_extract_exons.py). Reads were aligned to the genome using HISAT2 with default settings (version 2.1.0) (Kim *et al*. 2019). Samtools (version 1.7) (Li *et al*. 2009) was used to convert SAM files to BAM files, and to sort the BAM files for downstream analysis. Read counts were assembled using stringTie (version 1.3.5) (Pertea *et al*. 2015; Pertea *et al*. 2016), and FeatureCounts (version 1.6.0) (Liao *et al*. 2014). We used the accomPanying Python script from stringTie to prepare the output specifically for use in DESeq2 (Love *et al*. 2014).

### Differential exon usage analysis

Alternative splicing differences were determined by quantifying differential exon usage using the Bioconductor package DEXSeq (version 1.30.0) (Anders *et al*. 2013) in R. Expression profiles were built using FeatureCounts (Liao *et al*. 2014). Exons that showed differential expression with an adjusted *p* value false discovery rate less than 0.05 were considered differentially expressed in our analysis.

### Differential transcript expression analysis

We examined all cells versus muscle cells, and wild-type versus *etr-1(lq61)* muscle cells in separate analyses. Differential transcript expression was tested using the Bioconductor package DESeq2 (version 1.24.0) in R. Expression profiles were prepared using stringTie (Pertea *et al*. 2015; Pertea *et al*. 2016), and the accomPanying Python script was used to prepare a table for DESeq2. For both analyses, transcripts with a false-discovery rate adjusted *p*-value less than 0.05 were considered significantly differentially expressed. Preparation of tables and graphs was carried out in R (Wickham 2009).

### Gene Ontology analysis

Analysis was done using the Database for Annotation, Visualization and Integrated Discovery (DAVID: https://david.ncifcrf.gov/) (Huang da *et al*. 2009b; Huang da *et al*. 2009a). Alternative exon usage and alternative transcript expression were analyzed for biological process, cellular component, and molecular function separately. Gene lists were analyzed for terms that were significantly enriched at *q > 0.05* against all *C. elegans* genes. The complete GO term list for each separate test are in Supplementary Figures (3 and 5). Graphs were prepared in R (Wickham 2009).

### Sashimi plots

Sashimi plots were created using the Integrative Genomics Viewer (IGV), version 2.5.0 (Robinson *et al*. 2011; Thorvaldsdottir *et al*. 2013). Indexes to visualize the splice junctions on IGV were built using the samtools (version 1.7.0) index function (Li *et al*. 2009).

### RNA-mediated gene interference (RNAi)

RNAi was administered via feeding, following standard protocols and clones from the Source BioScience library (Nottingham UK). (Kamath and Ahringer 2003; Kamath *et al*. 2003) (Kamath et al., 2003). For each RNAi experiment we grew wild-type animals expressing *Pgcy-32::cfp* to visualize AQR and PQR on RNAi bacteria (Tamayo *et al*. 2013). For each independent set of RNAi experiments, *ceh-20(RNAi)* was used as a positive control, as RNAi of *ceh-20* results in robust AQR and PQR defects (Tamayo *et al*. 2013).

### Scoring AQR and PQR migration defects

AQR and PQR migration was scored using *gcy-32::cfp* as previously reported (Chapman *et al*. 2008; Sundararajan and Lundquist 2012; Ochs *et al*. 2020). Briefly, L4 animals were collected, mounted onto a 2% agar pad and immobilized by 5mM of NaN_3_. Five positions along the length of the animal were noted. Position 1 is the normal AQR position in the head just posterior to the posterior pharyngeal bulb; position 2 is posterior to the normal position but anterior to the vulva, position 3 is proximal to the vulva both anteriorly and posteriorly, position 4 is the normal birthplace of QR and QL, and position 5 is the normal location of PQR, just posterior to the anus. 100 animals were scored for each RNAi clone.

## Supporting information

R code for wild-type muscle to wild type all cells comparison

R code for wild type muscle to etr-1 muscle DEseq

R code for wildtype muscle to etr-1 muscle DEXseq

FASTQ file code for the project

## Acknowledgments

The authors thank the members of the Lundquist and Ackley labs for discussion, E. Struckhoff and Z. Grant for technical assistance, and R. Unckless for assistance and comments on the manuscript. This work was supported by National Institute of Health projects R21NS100483 to E.A.L. and D.M.M.; R01NS106951 and R01NS10054 to D.M.M.; and R01AI139154 to R.L.L. Some strains were provided by the CGC, which is funded by NIH Office of Research Infrastructure Programs (P40OD010440). Sequencing was conducted at the KU Genome Sequencing Core supported by the National Institute of General Medical Sciences (P20GM103638), and the Kansas Infrastructure Netrwork of Biomedical Research Excellence (P20GM103418) for computational support.

## References

1. Anders, S., D. J. McCarthy, Y. Chen, M. Okoniewski, G. K. Smyth. et al., 2013 Count-based differential expression analysis of RNA sequencing data using R and Bioconductor. Nat Protoc 8: 1765–1786.

2. Anyanful, A., Y. Sakube, K. Takuwa and H. Kagawa, 2001 The third and fourth tropomyosin isoforms of Caenorhabditis elegans are expressed in the pharynx and intestines and are essential for development and morphology. J Mol Biol 313: 525–537.

3. Banziger, C., D. Soldini, C. Schutt, P. Zipperlen, G. Hausmann et al., 2006 Wntless, a conserved membrane protein dedicated to the secretion of Wnt proteins from signaling cells. Cell 125: 509–522.

4. Berger, D. S., and A. N. Ladd, 2012 Repression of nuclear CELF activity can rescue CELF-regulated alternative splicing defects in skeletal muscle models of myotonic dystrophy. PLoS Curr 4: RRN1305.

5. Blech-Hermoni, Y., T. Dasgupta, R. J. Coram and A. N. Ladd, 2016 Identification of Targets of CUG-BP, Elav-Like Family Member 1 (CELF1) Regulation in Embryonic Heart Muscle. PLoS One 11: e0149061.

6. Boateng, R., and A. K. Allen, 2018 New Role for an Old Protein: An Educational Primer for Use with “The Identification of a Novel Mutant Allele of topoisomerase II in Caenorhabditis elegans Reveals a Unique Role in Chromosome Segregation During Spermatogenesis”. Genetics 208: 79–88.

7. Boateng, R., K. C. Q. Nguyen, D. H. Hall, A. Golden and A. K. Allen, 2017 Novel functions for the RNA-binding protein ETR-1 in Caenorhabditis elegans reproduction and engulfment of germline apoptotic cell corpses. Dev Biol 429: 306–320.

8. Broitman-Maduro, G., M. F. Maduro and J. H. Rothman, 2005 The noncanonical binding site of the MED-1 GATA factor defines differentially regulated target genes in the C. elegans mesendoderm. Dev Cell 8: 427–433.

9. Ch’ng, Q., L. Williams, Y. S. Lie, M. Sym, J. Whangbo et al., 2003 Identification of genes that regulate a left-right asymmetric neuronal migration in Caenorhabditis elegans. Genetics 164: 1355–1367.

10. Chalfie, M., and J. Sulston, 1981 Developmental genetics of the mechanosensory neurons of Caenorhabditis elegans. Dev Biol 82: 358–370.

11. Chapman, J. O., H. Li and E. A. Lundquist, 2008 The MIG-15 NIK kinase acts cell-autonomously in neuroblast polarization and migration in C. elegans. Dev Biol 324: 245–257.

12. Chen, L., Z. Liu, B. Zhou, C. Wei, Y. Zhou et al., 2016 CELF RNA binding proteins promote axon regeneration in C. elegans and mammals through alternative splicing of Syntaxins. Elife 5.

13. Chen, S., Y. Zhou, Y. Chen and J. Gu, 2018 fastp: an ultra-fast all-in-one FASTQ preprocessor. Bioinformatics 34: i884–i890.

14. Dasgupta, T., and A. N. Ladd, 2012 The importance of CELF control: molecular and biological roles of the CUG-BP, Elav-like family of RNA-binding proteins. Wiley Interdiscip Rev RNA 3: 104–121.

15. Daughters, R. S., D. L. Tuttle, W. Gao, Y. Ikeda, M. L. Moseley et al., 2009 RNA gain-of-function in spinocerebellar ataxia type 8. PLoS Genet 5: e1000600.

16. Ebbing, A., T. C. Middelkoop, M. C. Betist, E. Bodewes and H. C. Korswagen, 2019 Partially overlapping guidance pathways focus the activity of UNC-40/DCC along the anteroposterior axis of polarizing neuroblasts. Development 146.

17. Fox, R. M., J. D. Watson, S. E. Von Stetina, J. McDermott, T. M. Brodigan et al., 2007 The embryonic muscle transcriptome of Caenorhabditis elegans. Genome Biol 8: R188.

18. Fu, R., Y. Zhu, X. Jiang, Y. Li, M. Zhu et al., 2018 CCAR-1 affects hemidesmosome biogenesis by regulating unc-52/perlecan alternative splicing in the C. elegans epidermis. J Cell Sci 131.

19. Gomes, J. E., S. E. Encalada, K. A. Swan, C. A. Shelton, J. C. Carter et al., 2001 The maternal gene spn-4 encodes a predicted RRM protein required for mitotic spindle orientation and cell fate patterning in early C. elegans embryos. Development 128: 4301–4314.

20. Good, P. J., Q. Chen, S. J. Warner and D. C. Herring, 2000 A family of human RNA-binding proteins related to the Drosophila Bruno translational regulator. J Biol Chem 275: 28583–28592.

21. Harterink, M., D. H. Kim, T. C. Middelkoop, T. D. Doan, A. van Oudenaarden et al., 2011 Neuroblast migration along the anteroposterior axis of C. elegans is controlled by opposing gradients of Wnts and a secreted Frizzled-related protein. Development 138: 2915–2924.

22. Ho, T. H., D. Bundman, D. L. Armstrong and T. A. Cooper, 2005 Transgenic mice expressing CUG-BP1 reproduce splicing mis-regulation observed in myotonic dystrophy. Hum Mol Genet 14: 1539–1547.

23. Honigberg, L., and C. Kenyon, 2000 Establishment of left/right asymmetry in neuroblast migration by UNC-40/DCC, UNC-73/Trio and DPY-19 proteins in C. elegans. Development 127: 4655-4668.

24. Huang da, W., B. T. Sherman and R. A. Lempicki, 2009a Bioinformatics enrichment tools: paths toward the comprehensive functional analysis of large gene lists. Nucleic Acids Res 37: 1–13.

25. Huang da, W., B. T. Sherman and R. A. Lempicki, 2009b Systematic and integrative analysis of large gene lists using DAVID bioinformatics resources. Nat Protoc 4: 44–57.

26. Huang, X., P. Huang, M. K. Robinson, M. J. Stern and Y. Jin, 2003 UNC-71, a disintegrin and metalloprotease (ADAM) protein, regulates motor axon guidance and sex myoblast migration in C. elegans. Development 130: 3147–3161.

27. Josephson, M. P., R. Aliani, M. L. Norris, M. E. Ochs, M. Gujar et al., 2017 The Caenorhabditis elegans NF2/Merlin Molecule NFM-1 Nonautonomously Regulates Neuroblast Migration and Interacts Genetically with the Guidance Cue SLT-1/Slit. Genetics 205: 737–748.

28. Josephson, M. P., Y. Chai, G. Ou and E. A. Lundquist, 2016a EGL-20/Wnt and MAB-5/Hox Act Sequentially to Inhibit Anterior Migration of Neuroblasts in C. elegans. PLoS One 11: e0148658.

29. Josephson, M. P., A. M. Miltner and E. A. Lundquist, 2016b Nonautonomous Roles of MAB-5/Hox and the Secreted Basement Membrane Molecule SPON-1/F-Spondin in Caenorhabditis elegans Neuronal Migration. Genetics 203: 1747–1762.

30. Kagawa, H., K. Takuwa and Y. Sakube, 1997 Mutations and expressions of the tropomyosin gene and the troponin C gene of Caenorhabditis elegans. Cell Struct Funct 22: 213–218.

31. Kalsotra, A., X. Xiao, A. J. Ward, J. C. Castle, J. M. Johnson et al., 2008 A postnatal switch of CELF and MBNL proteins reprograms alternative splicing in the developing heart. Proc Natl Acad Sci U S A 105: 20333–20338.

32. Kamath, R. S., and J. Ahringer, 2003 Genome-wide RNAi screening in Caenorhabditis elegans. Methods 30: 313–321.

33. Kamath, R. S., A. G. Fraser, Y. Dong, G. Poulin, R. Durbin et al., 2003 Systematic functional analysis of the Caenorhabditis elegans genome using RNAi. Nature 421: 231–237.

34. Kim, D., J. M. Paggi, C. Park, C. Bennett and S. L. Salzberg, 2019 Graph-based genome alignment and genotyping with HISAT2 and HISAT-genotype. Nat Biotechnol 37: 907–915.

35. Kim, G., C. I. Pai, K. Sato, M. D. Person, A. Nakamura et al., 2015 Region-specific activation of oskar mRNA translation by inhibition of Bruno-mediated repression. PLoS Genet 11: e1004992.

36. Koorman, T., D. Klompstra, M. van der Voet, I. Lemmens, J. J. Ramalho et al., 2016 A combined binary interaction and phenotypic map of C. elegans cell polarity proteins. Nat Cell Biol 18: 337–346.

37. Kuroyanagi, H., Y. Watanabe and M. Hagiwara, 2013a CELF family RNA-binding protein UNC-75 regulates two sets of mutually exclusive exons of the unc-32 gene in neuron-specific manners in Caenorhabditis elegans. PLoS Genet 9: e1003337.

38. Kuroyanagi, H., Y. Watanabe, Y. Suzuki and M. Hagiwara, 2013b Position-dependent and neuron-specific splicing regulation by the CELF family RNA-binding protein UNC-75 in Caenorhabditis elegans. Nucleic Acids Res 41: 4015–4025.

39. Kuyumcu-Martinez, N. M., G. S. Wang and T. A. Cooper, 2007 Increased steady-state levels of CUGBP1 in myotonic dystrophy 1 are due to PKC-mediated hyperphosphorylation. Mol Cell 28: 68–78.

40. Ladd, A. N., N. Charlet and T. A. Cooper, 2001 The CELF family of RNA binding proteins is implicated in cell-specific and developmentally regulated alternative splicing. Mol Cell Biol 21: 1285–1296.

41. Ladd, A. N., G. Taffet, C. Hartley, D. L. Kearney and T. A. Cooper, 2005 Cardiac tissue-specific repression of CELF activity disrupts alternative splicing and causes cardiomyopathy. Mol Cell Biol 25: 6267–6278.

42. Lang, A. E., and E. A. Lundquist, 2021 The Collagens DPY-17 and SQT-3 Direct Anterior-Posterior Migration of the Q Neuroblasts in C. elegans. J Dev Biol 9.

43. Li, D., L. L. Bachinski and R. Roberts, 2001 Genomic organization and isoform-specific tissue expression of human NAPOR (CUGBP2) as a candidate gene for familial arrhythmogenic right ventricular dysplasia. Genomics 74: 396–401.

44. Li, H., B. Handsaker, A. Wysoker, T. Fennell, J. Ruan et al., 2009 The Sequence Alignment/Map format and SAMtools. Bioinformatics 25: 2078–2079.

45. Liao, Y., G. K. Smyth and W. Shi, 2014 featureCounts: an efficient general purpose program for assigning sequence reads to genomic features. Bioinformatics 30: 923–930.

46. Loria, P. M., A. Duke, J. B. Rand and O. Hobert, 2003 Two neuronal, nuclear-localized RNA binding proteins involved in synaptic transmission. Curr Biol 13: 1317–1323.

47. Love, M. I., W. Huber and S. Anders, 2014 Moderated estimation of fold change and dispersion for RNA-seq data with DESeq2. Genome Biol 15: 550.

48. Lundquist, E. A., R. K. Herman, T. M. Rogalski, G. P. Mullen, D. G. Moerman et al., 1996 The mec-8 gene of C. elegans encodes a protein with two RNA recognition motifs and regulates alternative splicing of unc-52 transcripts. Development 122: 1601–1610.

49. Maloof, J. N., J. Whangbo, J. M. Harris, G. D. Jongeward and C. Kenyon, 1999 A Wnt signaling pathway controls hox gene expression and neuroblast migration in C. elegans. Development 126: 37–49.

50. Masuda, H., K. Nakamura, N. Takata, B. Itoh, T. Hirose et al., 2012 MIG-13 controls anteroposterior cell migration by interacting with UNC-71/ADM-1 and SRC-1 in Caenorhabditis elegans. FEBS Lett 586: 740–746.

51. Merz, D. C., G. Alves, T. Kawano, H. Zheng and J. G. Culotti, 2003 UNC-52/perlecan affects gonadal leader cell migrations in C. elegans hermaphrodites through alterations in growth factor signaling. Dev Biol 256: 173–186.

52. Middelkoop, T. C., and H. C. Korswagen, 2014 Development and migration of the C. elegans Q neuroblasts and their descendants. WormBook: 1–23.

53. Middelkoop, T. C., L. Williams, P. T. Yang, J. Luchtenberg, M. C. Betist et al., 2012 The thrombospondin repeat containing protein MIG-21 controls a left-right asymmetric Wnt signaling response in migrating C. elegans neuroblasts. Dev Biol 361: 338–348.

54. Milne, C. A., and J. Hodgkin, 1999 ETR-1, a homologue of a protein linked to myotonic dystrophy, is essential for muscle development in Caenorhabditis elegans. Curr Biol 9: 1243–1246.

55. Mullen, G. P., T. M. Rogalski, J. A. Bush, P. R. Gorji and D. G. Moerman, 1999 Complex patterns of alternative splicing mediate the spatial and temporal distribution of perlecan/UNC-52 in Caenorhabditis elegans. Mol Biol Cell 10: 3205–3221.

56. Norris, A. D., S. Gao, M. L. Norris, D. Ray, A. K. Ramani et al., 2014 A pair of RNA-binding proteins controls networks of splicing events contributing to specialization of neural cell types. Mol Cell 54: 946–959.

57. Ochs, M. E., M. P. Josephson and E. A. Lundquist, 2020 The Predicted RNA-Binding Protein ETR-1/CELF1 Acts in Muscles To Regulate Neuroblast Migration in Caenorhabditis elegans. G3 (Bethesda) 10: 2365-2376.

58. Pan, C. L., P. D. Baum, M. Gu, E. M. Jorgensen, S. G. Clark et al., 2008 C. elegans AP-2 and retromer control Wnt signaling by regulating mig-14/Wntless. Dev Cell 14: 132–139.

59. Pertea, M., D. Kim, G. M. Pertea, J. T. Leek and S. L. Salzberg, 2016 Transcript-level expression analysis of RNA-seq experiments with HISAT, StringTie and Ballgown. Nat Protoc 11: 1650–1667.

60. Pertea, M., G. M. Pertea, C. M. Antonescu, T. C. Chang, J. T. Mendell et al., 2015 StringTie enables improved reconstruction of a transcriptome from RNA-seq reads. Nat Biotechnol 33: 290–295.

61. Robinson, J. T., H. Thorvaldsdottir, W. Winckler, M. Guttman, E. S. Lander et al., 2011 Integrative genomics viewer. Nat Biotechnol 29: 24–26.

62. Rogalski, T. M., E. J. Gilchrist, G. P. Mullen and D. G. Moerman, 1995 Mutations in the unc-52 gene responsible for body wall muscle defects in adult Caenorhabditis elegans are located in alternatively spliced exons. Genetics 139: 159–169.

63. Rogalski, T. M., G. P. Mullen, J. A. Bush, E. J. Gilchrist and D. G. Moerman, 2001 UNC-52/perlecan isoform diversity and function in Caenorhabditis elegans. Biochem Soc Trans 29: 171–176.

64. Rogalski, T. M., B. D. Williams, G. P. Mullen and D. G. Moerman, 1993 Products of the unc-52 gene in Caenorhabditis elegans are homologous to the core protein of the mammalian basement membrane heparan sulfate proteoglycan. Genes Dev 7: 1471–1484.

65. Santoro, M., A. Modoni, M. Masciullo, T. Gidaro, A. Broccolini et al., 2010 Analysis of MTMR1 expression and correlation with muscle pathological features in juvenile/adult onset myotonic dystrophy type 1 (DM1) and in myotonic dystrophy type 2 (DM2). Exp Mol Pathol 89: 158–168.

66. Savkur, R. S., A. V. Philips and T. A. Cooper, 2001 Aberrant regulation of insulin receptor alternative splicing is associated with insulin resistance in myotonic dystrophy. Nat Genet 29: 40–47.

67. Sawa, H., and H. C. Korswagen, 2013 Wnt signaling in C. elegans. WormBook: 1–30.

68. Schoser, B., and L. TimChenko, 2010 Myotonic dystrophies 1 and 2: complex diseases with complex mechanisms. Curr Genomics 11: 77–90.

69. Singh, N. N., J. Seo, E. W. Ottesen, M. Shishimorova, D. Bhattacharya et al., 2011 TIA1 prevents skipping of a critical exon associated with spinal muscular atrophy. Mol Cell Biol 31: 935–954.

70. Sofola, O. A., P. Jin, Y. Qin, R. Duan, H. Liu et al., 2007 RNA-binding proteins hnRNP A2/B1 and CUGBP1 suppress fragile X CGG premutation repeat-induced neurodegeneration in a Drosophila model of FXTAS. Neuron 55: 565–571.

71. Spencer, W. C., R. McWhirter, T. Miller, P. Strasbourger, O. Thompson et al., 2014 Isolation of specific neurons from C. elegans larvae for gene expression profiling. PLoS One 9: e112102.

72. Spike, C. A., A. G. Davies, J. E. Shaw and R. K. Herman, 2002 MEC-8 regulates alternative splicing of unc-52 transcripts in C. elegans hypodermal cells. Development 129: 4999–5008.

73. Sulston, J. E., and H. R. Horvitz, 1977 Post-embryonic cell lineages of the nematode, Caenorhabditis elegans. Dev Biol 56: 110–156.

74. Sundararajan, L., and E. A. Lundquist, 2012 Transmembrane proteins UNC-40/DCC, PTP-3/LAR, and MIG-21 control anterior-posterior neuroblast migration with left-right functional asymmetry in Caenorhabditis elegans. Genetics 192: 1373-1388.

75. Sundararajan, L., M. L. Norris and E. A. Lundquist, 2015 SDN-1/Syndecan Acts in Parallel to the Transmembrane Molecule MIG-13 to Promote Anterior Neuroblast Migration. G3 (Bethesda).

76. Sundararajan, L., M. L. Norris, S. Schoneich, B. D. Ackley and E. A. Lundquist, 2014 The fat-like cadherin CDH-4 acts cell-non-autonomously in anterior-posterior neuroblast migration. Dev Biol 392: 141–152.

77. Sureau, A., J. Sauliere, A. Expert-Bezancon and J. Marie, 2011 CELF and PTB proteins modulate the inclusion of the beta-tropomyosin exon 6B during myogenic differentiation. Exp Cell Res 317: 94–106.

78. Suzuki, H., Y. Jin, H. Otani, K. Yasuda and K. Inoue, 2002 Regulation of alternative splicing of alpha-actinin transcript by Bruno-like proteins. Genes Cells 7: 133–141.

79. Tamayo, J. V., M. Gujar, S. J. Macdonald and E. A. Lundquist, 2013 Functional transcriptomic analysis of the role of MAB-5/Hox in Q neuroblast migration in Caenorhabditis elegans. BMC Genomics 14: 304.

80. Taylor, S. R., G. Santpere, A. Weinreb, A. Barrett, M. B. Reilly et al., 2020 Molecular topography of an entire nervous system. bioRxiv: 2020.2012.2015.422897.

81. Terenzi, F., K. R. Brimacombe, M. S. Penn and A. N. Ladd, 2009 CELF-mediated alternative splicing is required for cardiac function during early, but not later, postnatal life. J Mol Cell Cardiol 46: 395–404.

82. Thorvaldsdottir, H., J. T. Robinson and J. P. Mesirov, 2013 Integrative Genomics Viewer (IGV): high-performance genomics data visualization and exploration. Brief Bioinform 14: 178–192.

83. TimChenko, N. A., Z. J. Cai, A. L. Welm, S. Reddy, T. Ashizawa et al., 2001 RNA CUG repeats sequester CUGBP1 and alter protein levels and activity of CUGBP1. J Biol Chem 276: 7820–7826.

84. TimChenko, N. A., R. Patel, P. Iakova, Z. J. Cai, L. Quan et al., 2004 Overexpression of CUG triplet repeat-binding protein, CUGBP1, in mice inhibits myogenesis. J Biol Chem 279: 13129-13139.

85. Wang, X., J. Liu, Z. Zhu and G. Ou, 2015 The heparan sulfate-modifying enzyme glucuronyl C5-epimerase HSE-5 controls Caenorhabditis elegans Q neuroblast polarization during migration. Dev Biol.

86. Watabe, E., S. Ono and H. Kuroyanagi, 2018 Alternative splicing of the Caenorhabditis elegans lev-11 tropomyosin gene is regulated in a tissue-specific manner. Cytoskeleton (Hoboken) 75: 427–436.

87. Wickham, H., 2009 ggplot2: Elegant Graphics for Data Analysis. Springer-Verlag New York.

88. Wijsman, E. M., N. D. Pankratz, Y. Choi, J. H. Rothstein, K. M. Faber et al., 2011 Genome-wide association of familial late-onset Alzheimer’s disease replicates BIN1 and CLU and nominates CUGBP2 in interaction with APOE. PLoS Genet 7: e1001308.

89. Williams, B. D., and R. H. Waterston, 1994 Genes critical for muscle development and function in Caenorhabditis elegans identified through lethal mutations. J Cell Biol 124: 475–490.

90. Zinovyeva, A. Y., Y. Yamamoto, H. Sawa and W. C. Forrester, 2008 Complex Network of Wnt Signaling Regulates Neuronal Migrations During Caenorhabditis elegans Development. Genetics 179: 1357–1371.

