## Supplemental material legend for "*Caenorhabditis elegans* ETR-1/CELF has broad effects on the muscle cell transcriptome, including genes that regulate translation and neuroblast migration"

Code files and their descriptions:

File name: "etr-1_project_code_publication"
This file contains all of the work done on the Unix command line.
That includes: quality control of the reads, cleaning up the reads, aligning the reads, converting to BAM format, transcriptome
building and preparing for downstream analysis in R.

File name: "wildType_etr1_exon_usage.R"
This file contains the analysis done in R. This file was conducting the differential exon usage comparison, comparing wild-type
muscle to etr-1 muscle. The main package used in this file was DEXSeq

File name: "wildType_etr1_muscle_transcript.R"
This file contains the differential transcript comparison of wild-type muscle to etr-1 muscle. The transcriptome was built using
stringTie and the downstream analysis was carried out using the package DESeq2.

File name: "wildType_muscle_whole_gene.R"
This file contains the differential gene expression analysis when comparing wild-type muscle to wild-type whole animal. This analysis
was conducted in R using the package DESeq2.
