## Supplementary material for "*Caenorhabditis elegans* ETR-1/CELF has broad effects on the muscle cell transcriptome, including genes that regulate translation and neuroblast migration": R code for wild-type muscle to wild type all cells comparison

library(DESeq2)

library(tidyverse)

library(hexbin)

library(pheatmap)

library(RColorBrewer)

library(ggrepel)

library(apeglm)

library(ashr)

library(reshape)

library(rtracklayer)

library(org.Ce.eg.db)

library(EnhancedVolcano)

genes <- read.csv("stringTie_genes.csv", header = TRUE);

head(genes)

fileNames <- colnames(genes %>%

dplyr::select(contains("RM")))

fileNames

sampleNames <- fileNames

sampleNames <- regmatches(sampleNames, regexpr("RM.\\d\\d", sampleNames))

sampleNames

sampleNamesRename <- gsub(pattern = "[.]",

replacement = "_",

sampleNames);

sampleNamesRename

colnames(genes)[2:13] <- sampleNamesRename

head(genes)

coldata <- data.frame(sampleNamesRename,

genotype = c("wt", "etr1", "etr1", "wt", "etr1", "wt", "etr1", "etr1", "wt", "etr1", "wt", "wt"),

tissue = rep(c("whole", "muscle"), each = 6))

coldata

coldata2 <- coldata[, -1]

coldata2

rownames(coldata2) <- sampleNamesRename

coldata2

#Setting up for muscle cell comparision to whole animal wildtpye

coldata

coldataWildType <- coldata %>%

filter(genotype == "wt") %>%

dplyr::select(genotype, tissue)

rownames(coldataWildType) <- sampleNamesRename[c(1, 4, 6, 9, 11, 12)]

coldataWildType

genesWildType <- genes %>%

dplyr::select(c(2, 5, 7, 10, 12, 13))

rownames(genesWildType) <- genes$gene_id

head(genesWildType)

ddsWildType <- DESeqDataSetFromMatrix(countData = genesWildType,

colData = coldataWildType,

design = ~ tissue)

ddsWildType

ddsWildType$tissue <- relevel(ddsWildType$tissue, ref = "whole")

ddsWildType <- estimateSizeFactors(ddsWildType)

ddsWildType <- estimateDispersions(ddsWildType)

plotDispEsts(ddsWildType)

ddsWildType <- DESeq(ddsWildType)

resWildType <- results(ddsWildType, alpha = 0.05)

summary(resWildType)

resWildType <- resWildType[order(resWildType$padj, decreasing = FALSE, na.last = NA), ]

resWildType$symbol <- mapIds(org.Ce.eg.db,

keys = rownames(resWildType),

column = "SYMBOL",

keytype = "WORMBASE",

multiVals = "first")

padj.cutoff <- 0.05

lfc.cutoff <- 2

resWT_tb <- resWildType %>%

data.frame() %>%

rownames_to_column(var = "WB_GeneID") %>%

as_tibble()

head(resWT_tb)

sigWT <- resWT_tb %>%

filter(padj < padj.cutoff & abs(log2FoldChange) > lfc.cutoff)

head(sigWT)

EnhancedVolcano(resWildType,

x = 'log2FoldChange',

y = 'padj',

lab = rownames(resWildType),

ylab = bquote(~-Log[10]~adjusted~italic(P)),

selectLab = padj.cutoff,

ylim = c(0,100),

pCutoff = 0.05,

FCcutoff = 1.0,

title = NULL,

subtitle = NULL,

legendVisible = FALSE)

write.csv(resWT_tb,

file = "wildType_muscle_whole_gene.csv")
