## Supplementary material for "*Caenorhabditis elegans* ETR-1/CELF has broad effects on the muscle cell transcriptome, including genes that regulate translation and neuroblast migration": R code for wild type muscle to etr-1 muscle DEseq

library(DESeq2)

library(tidyverse)

library(ggplot2)

library(hexbin)

library(pheatmap)

library(RColorBrewer)

library(ggrepel)

library(apeglm)

library(ashr)

library(reshape)

library(rtracklayer)

library(org.Ce.eg.db)

library(EnhancedVolcano)

transcripts <- read.csv("stringTie_transcripts.csv", header = TRUE)

head(transcripts)

fileNames <- colnames(genes %>%

dplyr::select(contains("RM")))

sampleNames <- fileNames

sampleNames <- regmatches(sampleNames, regexpr("RM.\\d\\d", sampleNames))

sampleNamesRename <- gsub(pattern = "[.]",

replacement = "_",

sampleNames);

coldata <- data.frame(sampleNamesRename,

genotype = c("wt", "etr1", "etr1", "wt", "etr1", "wt", "etr1", "etr1", "wt", "etr1", "wt", "wt"),

tissue = rep(c("whole", "muscle"), each = 6))

coldata2 <- coldata[, -1]

rownames(coldata2) <- sampleNamesRename

colnames(transcripts)[2:13] <- sampleNamesRename

head(transcripts)

#Setting up colData for wildtype to mutant muscle transcripts

colDataMuscle <- coldata %>%

dplyr::filter(tissue == "muscle")

rownames(colDataMuscle) <- colDataMuscle$sampleNamesRename

colDataMuscle <- colDataMuscle %>%

dplyr::select(genotype, tissue)

transMuscle <- transcripts %>%

dplyr::select(rownames(colDataMuscle))

rownames(transMuscle) <- transcripts$transcript_id

head(transMuscle)

##Entering data into a DESeq2 object##

#Entering in data into DESeqDataSet

ddsMuscle <- DESeqDataSetFromMatrix(countData = transMuscle,

colData = colDataMuscle,

design = ~genotype)

ddsMuscle

ddsMuscle$genotype <- relevel(ddsMuscle$genotype, ref = "wt")

ddsMuscle <- estimateSizeFactors(ddsMuscle)

ddsMuscle <- DESeq2::estimateDispersions(ddsMuscle, fitType = "local")

plotDispEsts(ddsMuscle)

ddsMuscle <- DESeq(ddsMuscle)

resMuscle <- results(ddsMuscle, alpha = 0.05)

summary(resMuscle)

resMuscleOrder <- resMuscle[order(resMuscle$padj, decreasing = FALSE, na.last = NA), ]

resMuscleDF <- data.frame(resMuscleOrder)

resMuscleDF$transcript_id <- rownames(resMuscleDF)

gtf <- rtracklayer::import("ce.gtf")

gtf_df <- data.frame(gtf)

rm(gtf)

gtf_df <- gtf_df %>%

dplyr::select(gene_id, gene_name, transcript_id)

head(gtf_df)

resMuscleTrans <- merge(gtf_df, resMuscleDF, by = "transcript_id")

resMuscleTrans <- distinct(resMuscleTrans)

resMuscleTrans <- resMuscleTrans[order(resMuscleTrans$padj, decreasing = FALSE, na.last = NA), ]

head(resMuscleTrans)

sum(resMuscleTrans$padj < 0.05)

EnhancedVolcano(resMuscle,

lab = 'transcript_id',

x = 'log2FoldChange',

y = 'padj',

title = NULL,

subtitle = NULL,

selectLab = "WBGene00001340",

pCutoff = 0.05,

FCcutoff = 1,

ylim = c(0, 20),

xlim = c(-20, 20),

legendVisible = FALSE)

padj.cutoff <- 0.05

lfc.cutoff <- 1

resMuscleTrans <- resMuscleTrans %>%

filter(padj < padj.cutoff & abs(log2FoldChange) > lfc.cutoff)

write.csv(resMuscleTrans, file = "wildType_mutant_muscle_comp_transcript.csv")
