## Supplementary material for "*Caenorhabditis elegans* ETR-1/CELF has broad effects on the muscle cell transcriptome, including genes that regulate translation and neuroblast migration": R code for wildtype muscle to etr-1 muscle DEXseq

library(DEXSeq)

library(tidyverse)

library(BiocParallel)

library(EnhancedVolcano)

source("load_SubreadOutput.R")

fileNames <- list.files(pattern = "RM-\\d\\d")

grep("RM-\\d\\d_etr1", fileNames, value = TRUE)

sampleNames <- fileNames[7:12]

sampleNames <- regmatches(sampleNames, regexpr("RM-\\d\\d", sampleNames))

sampleNamesRename <- gsub(pattern = "-",

replacement = "_",

sampleNames);

muscle <- read.table("wildType_mutant_muscle_featureCounts.txt", header = TRUE)

muscle <- muscle %>%

dplyr::select(1, 7:12)

colnames(muscle)[2:7] <- rownames(muscleSamp)

muscleSamp <- data.frame(row.names = c("RM_70", "RM_75", "RM_89", "RM_84", "RM_91", "RM_94"),

condition = c("etr1", "etr1", "etr1", "wt", "wt", "wt"))

muscleSamp$condition <- relevel(muscleSamp$condition, ref = "wt")

dxdMuscle <- DEXSeqDataSetFromFeatureCounts(countfile = "wildType_mutant_muscle_featureCounts.txt",

flattenedfile = "flat_ce.gtf",

sampleData = muscleSamp,

design = ~ sample + exon + condition:exon)

BPPARAM <- MulticoreParam(4)

dxdMuscle <- estimateSizeFactors(dxdMuscle)

dxdMuscle <- estimateDispersions(dxdMuscle, BPPARAM = BPPARAM)

plotDispEsts(dxdMuscle)

dxdMuscle <- testForDEU(dxdMuscle, BPPARAM = BPPARAM)

dxdMuscle <- estimateExonFoldChanges(dxdMuscle, fitExpToVar="condition", BPPARAM = BPPARAM)

resMuscle <- DEXSeqResults(dxdMuscle)
