## Supplementary material for "*Caenorhabditis elegans* ETR-1/CELF has broad effects on the muscle cell transcriptome, including genes that regulate translation and neuroblast migration": FASTQ file code for the project

##########Quality control with FASTQC##########

fastqc $FASTQ_DIR/RM-70_etr1_R1_002.fastq.gz \

$FASTQ_DIR/RM-70_etr1_R2_002.fastq.gz \

$FASTQ_DIR/RM-75_etr1_R1_002.fastq.gz \

$FASTQ_DIR/RM-75_etr1_R2_002.fastq.gz \

$FASTQ_DIR/RM-84_wt_R1_002.fastq.gz \

$FASTQ_DIR/RM-84_wt_R2_002.fastq.gz \

$FASTQ_DIR/RM-89_etr1_R1_002.fastq.gz \

$FASTQ_DIR/RM-89_etr1_R2_002.fastq.gz \

$FASTQ_DIR/RM-91_wt_R1_002.fastq.gz \

$FASTQ_DIR/RM-91_wt_R2_002.fastq.gz \

$FASTQ_DIR/RM-94_wt_R1_002.fastq.gz \

$FASTQ_DIR/RM-94_wt_R2_002.fastq.gz \

--outdir $FASTQC_DIR \

--contaminants /home/mattochs/RNA_Seq_Tools/illumina_contaminant_list.txt

fastqc $FASTQ_DIR/RM-02_wt_R1_002.fastq.gz \

$FASTQ_DIR/RM-02_wt_R2_002.fastq.gz \

$FASTQ_DIR/RM-03_etr1_R1_002.fastq.gz \

$FASTQ_DIR/RM-03_etr1_R2_002.fastq.gz \

$FASTQ_DIR/RM-04_etr1_R1_002.fastq.gz \

$FASTQ_DIR/RM-04_etr1_R2_002.fastq.gz \

$FASTQ_DIR/RM-05_wt_R1_002.fastq.gz \

$FASTQ_DIR/RM-05_wt_R2_002.fastq.gz \

$FASTQ_DIR/RM-06_etr1_R1_002.fastq.gz \

$FASTQ_DIR/RM-06_etr1_R2_002.fastq.gz \

$FASTQ_DIR/RM-07_wt_R1_002.fastq.gz \

$FASTQ_DIR/RM-07_wt_R2_002.fastq.gz \

--outdir $FASTQC_DIR \

--contaminants /home/mattochs/RNA_Seq_Tools/illumina_contaminant_list.txt
